# Data-Driven Discovery of Mechanistic Ecosystem Models with LLMs

**DOI:** 10.1101/2025.07.14.664628

**Authors:** Scott Spillias, Jacob Rogers, Fabio Boschetti, Beth Fulton, Magda Guglielmo, SukYee Yong, Rowan Trebilco

**Affiliations:** CSIRO Environment, Hobart, Australia; Centre for Marine Socio-Ecology, University of Tasmania, Hobart, Australia; CSIRO Environment, St. Lucia, Australia; CSIRO Environment, IOMRC Crawley, Australia; CSIRO IM&T, Eveleigh, Australia

**Keywords:** Artificial Intelligence, Ecological modelling, Evolutionary Algorithms, Large Language Models, Marine Ecosystems, Crown-of-Thorns Starfish, Great Barrier Reef, time-series forecasting

## Abstract

Ecosystem models are essential for ecosystem management, but their development traditionally requires significant time and expertise, creating bottlenecks in addressing urgent environmental challenges.

We present ***LEMMA*** (LLM Enabled Mechanistic Modelling for ecosystem Assessment), a framework that programmatically generates and iteratively refines mechanistic ecosystem models by combining large language models (LLMs) for equation synthesis and parameter search, evolutionary algorithms for structural optimization, and Template Model Builder (TMB) for efficient parameter estimation.

We critically review *LEMMA*’s ability to recover known ecological relationships through two complementary marine case studies: (1) a nutrient-phytoplankton-zooplankton model, and (2) a Crown-of-Thorns starfish (COTS) model. In the first case, our best models displayed almost perfect recovery of known ecological dynamics while maintaining strong predictive performance across multivariate time-series. In the second case, best *LEMMA* generated models approached human expert models in terms of their ability to successfully capture COTS outbreak dynamics and demonstrated strong out-of-sample predictive power.

*LEMMA* produces interpretable models with meaningful parameters that capture real biological processes, facilitating scientific insight and potentially accelerating management applications. By dramatically accelerating model development while offering ecological interpretability, *LEMMA* offers a powerful new tool for addressing urgent ecological challenges in a changing world.

## 1 Introduction

Ecosystem models provide invaluable information for managing complex interactions between nature and people (McCarthy and Possingham, 2004; Holden and Ellner, 2016), but their development traditionally requires significant time and expertise, creating a bottleneck in addressing urgent environmental challenges (Dichmont et al., 2017; Holden et al., 2024), particularly as climate change demands rapid, adaptable approaches for ecosystem management (Weiskopf et al., 2020; Malhi et al., 2020).

Artificial Intelligence (AI) offers great promise as a part of the solution to these modelling challenges, with potential to accelerate model development and enhance adaptability (Spillias et al., 2024a). While initial efforts to apply AI in ecological modelling focused on machine learning approaches that rely on black-box methods (Morales-García et al., 2024), emerging techniques in equation discovery and automated scientific discovery show particular promise (Huntingford et al., 2024; Floryan and Graham, 2022). These methods can derive interpretable mathematical relationships directly from data, offering advantages over statistical emulators when modelling novel environmental conditions (Schaeffer, 2017; Chen et al., 2024; Karniadakis et al., 2021).

A recent development in AI is the rise of large language models (LLMs): neural networks trained on vast text corpora, including scientific literature, code, and web content. LLMs can generate and interpret language, code, and equations, encoding broad domain knowledge that can be leveraged for scientific modelling tasks (Mammides and Papadopoulos, 2024; Wills et al., 2024). Attempts to leverage LLMs for direct time-series prediction (Zhang et al., 2024; Su et al., 2024; Hassani and Silva, 2024; Gandhi et al., 2024; Bylund and Johansson, 2024; Cao et al., 2023; Li et al., 2024; Garza et al., 2023), though successful in other fields, are unsuited for producing reliable ecological insights or testing management interventions. For instance, work on multimodal LLMs for environmental prediction (Li et al., 2024) achieves impressive accuracy in forecasting physical variables like streamflow and water temperature, but does not address the mechanistic relationships needed for ecosystem management and applications.

Rather than using AI to replace traditional modelling approaches, recent advances in AI coding capabilities suggest a more promising direction (Xu et al., 2021). Recent demonstrations of LLMs automating scientific processes, from autonomous chemical experimentation (Burger et al., 2023) to biomedical research (Wang et al., 2024), evidence synthesis (Spillias et al., 2024b) and even fully automated scientific discovery (Kramer et al., 2023), highlight their potential for systematic scientific work. The key challenge lies in developing frameworks that can systematically harness these capabilities while ensuring scientific rigor and maintaining human oversight in the discovery process (Kramer et al., 2023; Spillias et al., 2024a). To the best of our knowledge, such an approach has not been attempted yet in ecological modelling.

To address this challenge, we present “LLM-Enabled Mechanistic Modelling for ecosystem Assessment” (LEMMA), a novel framework that requires minimal inputs, only time-series data and research questions, and aims to produce ecologically sound mathematical models that explain the time-series data (Figure 1). LEMMA addresses the inverse problem of inferring ecologically meaningful mechanistic models and parameters that causally explain observed data. The inferred models can then be used to test management interventions and scenarios. When solving inverse problems through numerical optimization, practitioners usually predetermine the model and its allowable parameter ranges, which defines both the parameter space and its mapping to the observation space. Less commonly, the model itself is inferred directly from observations, as in Scientific Machine Learning (SciML) (Willard et al., 2022), System Identification (Chiuso and Pillonetto, 2019), and Automated Algorithm Discovery (Blazek et al., 2024). Successful model reconstruction in such scenarios requires extensive data to ensure statistical relations are represented in the observations. In addition, a common challenge across all inverse problem approaches is the need to impose constraints that prevent numerically accurate yet empirically unrealistic solutions.

**Figure 1:**
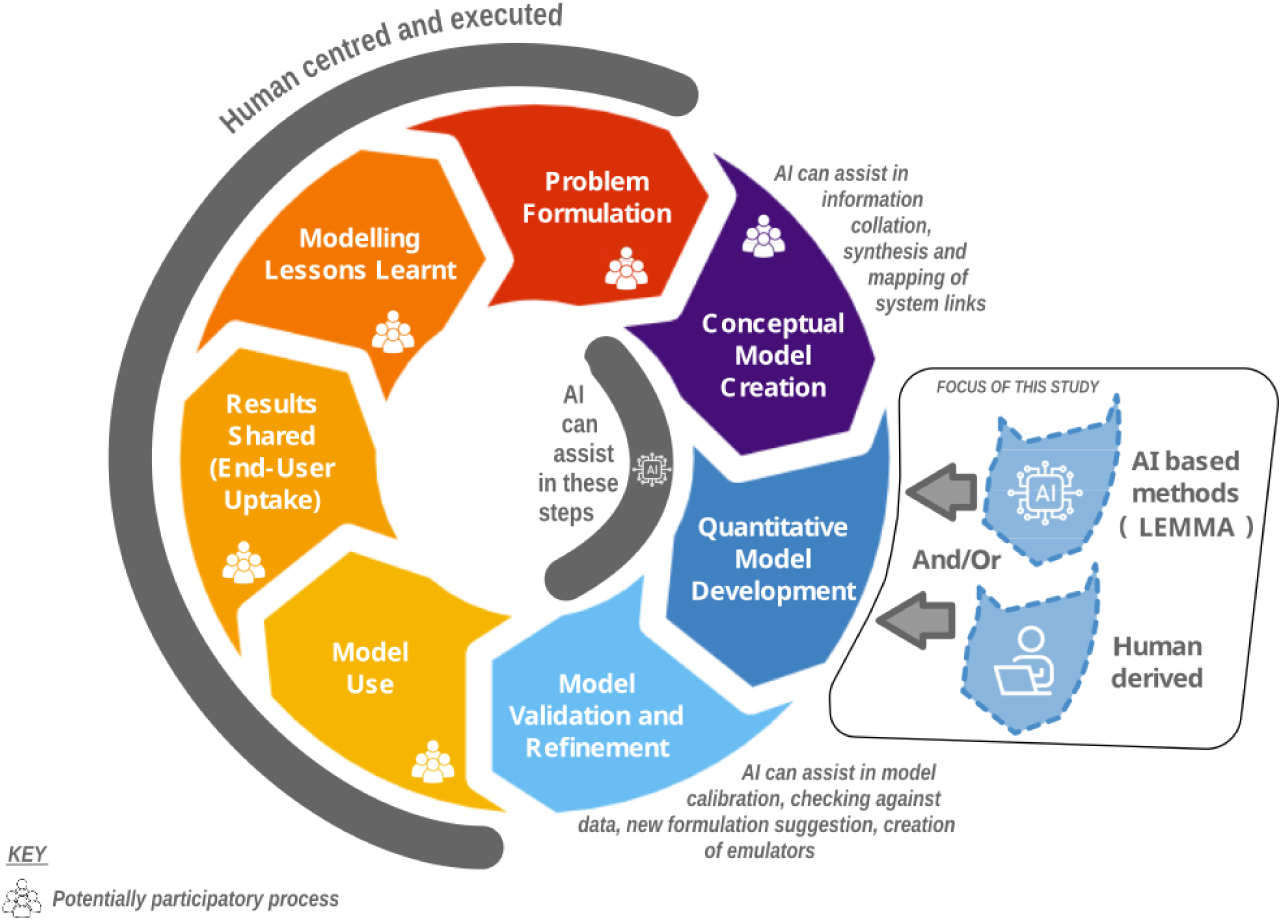
Stylised representation of the iterative modelling process that LEMMA aims to support. Whilst human experts drive the majority of the process, we show that AI-driven processes could play an important role in the Model Development stage.

**Figure 2:**
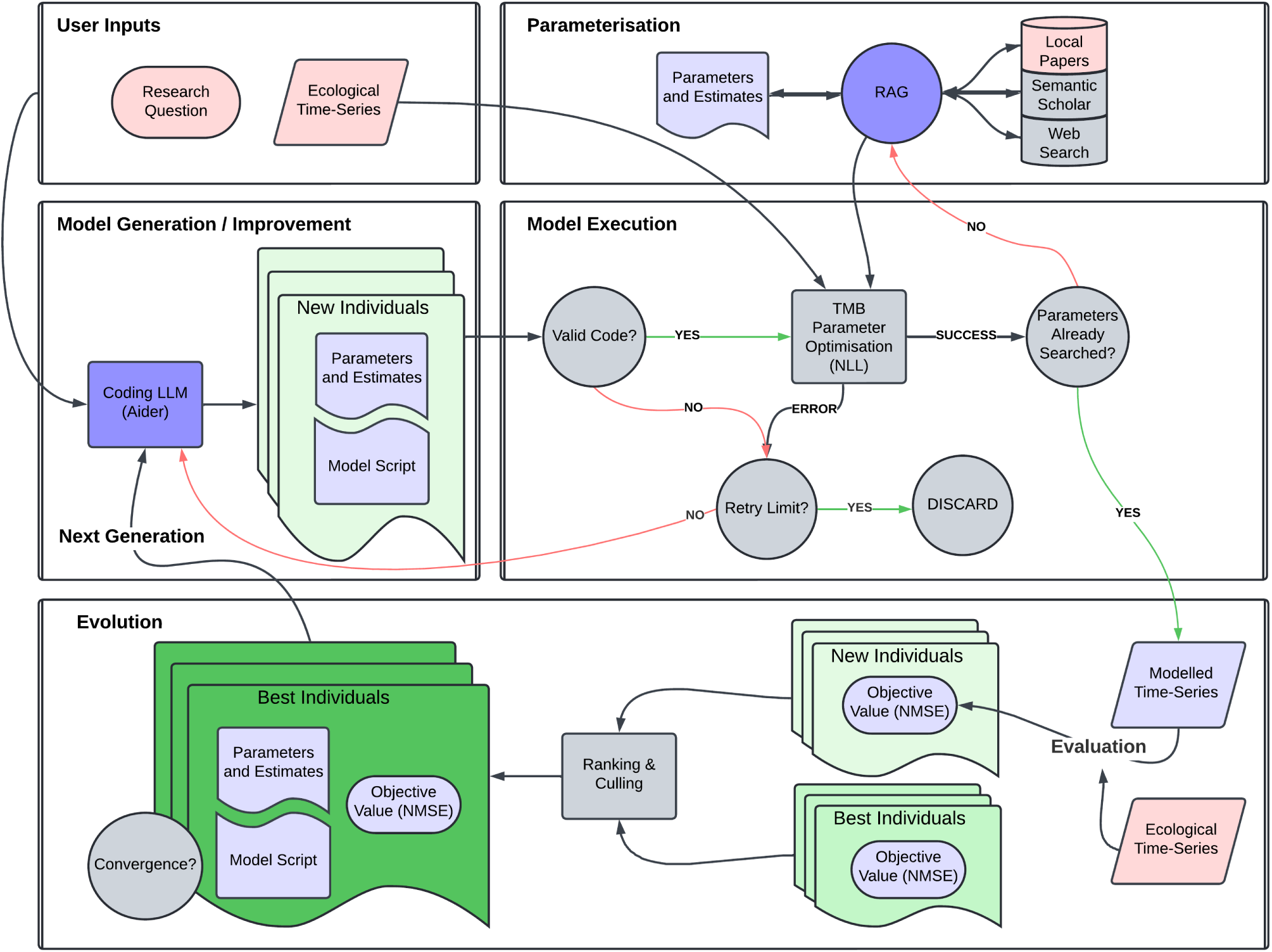
Conceptual diagram of the automated ecological modelling framework, LEMMA. The workflow consists of five main components: (1) User Inputs, where research questions and ecological time-series data are provided; (2) Parameterisation, utilizing RAG-enhanced literature search to estimate parameter values; (3) Model Generation/Improvement, where the Coding LLM creates new individuals with model scripts and parameters; (4) Model Execution, where the LLM’s model code is implemented and TMB is used to optimise parameter values; and (5) Evolution, which evaluates model performance through individual assessment, error handling, and ranking-based

LEMMA differs from traditional and SciML approaches by leveraging the broad scientific and ecological knowledge encoded in LLMs from their pre-training on diverse sources such as peer-reviewed literature, textbooks, and code repositories. This embedded knowledge is used to impose ecologically meaningful constraints on both model structures and parameter ranges. In addition, LEMMA incorporates a Retrieval-Augmented Generation (RAG) process to identify parameter values and plausible bounds from trusted local repositories and online scientific sources, ensuring biologically realistic parameterization. Within this framework, LEMMA employs Template Model Builder (TMB; an R/C++ library for efficient parameter estimation in complex nonlinear models) as its computational backbone, providing a rigorous statistical foundation for ecological modelling. The system operates iteratively: an LLM proposes candidate model structures as TMB-compatible equations, which are evaluated against time-series data using normalized objective functions (where lower values indicate better fit). These models then undergo evolutionary optimization across multiple generations, with high-performing structures (“individuals”) retained and refined while under-performing models are discarded. This evolutionary process systematically yields increasingly accurate representations of ecosystem dynamics and enables exploration of alternative mechanistic hypotheses with quantified support.

We test LEMMA’s ability to construct plausible ecological models through two complementary case studies that test different aspects of ecological modelling, each involving both dependent variables (state variables predicted by the model) and forcing variables (external drivers affecting the system). First, we evaluate the framework’s ability to recover fundamental ecological understanding using synthetic data generated from a well-established nutrient-phytoplankton-zooplankton (NPZ) model (Edwards and Brindley, 1999). This experiment tests LEMMA’s equation-learning capabilities by comparing discovered equations against known mathematical relationships that represent core ecological processes. For our NPZ case study, we additionally evaluate models using ecological accuracy scores (on a scale of 0-8), which measure how well the generated models recover known ecological mechanisms from the reference model. These scores assess specific components like nutrient uptake, phytoplankton growth, and zooplankton dynamics, with higher scores indicating closer alignment with established ecological understanding. Second, we assess LEMMA’s ability to provide management-relevant predictions using synthetic data based on Crown-of-Thorns starfish (COTS) populations on the Great Barrier Reef, derived from existing MICE models (Morello et al., 2014; Rogers and Plagányi, 2022; Plagányi et al., 2014; Condie et al., 2021). This case study included three dependent variables (COTS abundance, fast-growing coral cover, and slow-growing coral cover) and two forcing variables (temperature and larval immigration), testing the framework’s robustness in a complex predator-prey system. We implemented a factorial design comparing three state-of-the-art LLMs, GPT-5, Claude-Sonnet-4.5, and Gemini-2.5-Pro, across both case studies, with multiple replicates and evolutionary runs. This comparative approach allowed us to evaluate differences in convergence, predictive accuracy, and ecological realism, demonstrating that LEMMA can both rediscover theoretical relationships and approximate expert-level performance.

## 2 Methods

At its core, LEMMA integrates LLMs for generating and modifying model structures, Template Model Builder (TMB) for statistical parameter estimation, and evolutionary algorithms for systematic model improvement. All of the code and data underpinning this study are available at the Github repository: https://github.com/s-spillias/EMs-with-LLMs/.

### 2.1 LEMMA Framework

#### 2.1.1 Model Generation and Improvement

LEMMA uses LLMs to write and modify computer code through Aider (Gauthier, 2024), which is a coding assistant that can create, modify, and interpret local files. Aider can be used in the command-line or called within python scripts, as we have done here, and can receive text and/or images as input, depending on whether the underlying LLM is ‘multi-modal’ (i.e., can interpret text and images). Each model instance, referred to as an individual in our evolutionary framework, has two fundamental ‘chromosomes’: a TMB-compatible dynamic model written in C++ that implements a system of equations (model.cpp) and (2) a parameters file containing initial values and bounds (parameters.json). Both of these files are provided to the LLM (via Aider) at each initialization and modification step. We also ask the LLM to provide a third documentation file explaining the ecological meaning of the proposed actions for the benefit of the user and to audit the model development process (intention.txt) (see Supplement for the complete prompt to Aider).

The LLM generates initial parameter estimates for pre-testing model structure before optimization begins. For each parameter, it assigns a priority number that determines optimization order, following established practices in ecosystem modelling (Plagányi et al., 2014).

During non-initial steps, if multi-modal (i.e. can receive images as input) the LLM analyzes performance plots comparing predictions to historical data, otherwise the LLM receives a structured file showing the model fit residuals. After interpreting the model fit, LEMMA makes targeted, ecologically meaningful changes to model equations, implementing one modification at a time to maintain transparency and traceability of successful modelling strategies (see Supplement).

#### 2.1.2 User Inputs

The LEMMA framework requires only minimal user input to initiate the modelling process. At the outset, the user provides a short natural language description of the ecological system and research question, which guides LEMMA in generating ecologically relevant model structures. Two timeseries data files are also supplied: a response file containing the dependent variables (state variables) to be predicted by the model, and a forcing file containing external drivers that influence system dynamics. Column headers in these files define the variables that are directly observed and available for model fitting. Importantly, the framework is not restricted to these variables; the LEMMA may introduce additional latent variables or intermediate processes when such constructs are ecologically justified and improve model performance. This flexibility allows LEMMA to explore a broader range of mechanistic hypotheses than would be possible if limited to observed variables alone.

In addition to these core inputs, users may optionally specify configuration parameters that control the evolutionary process and LLM behaviour, including the sampling temperature, the number of individuals per generation, the number of generations, the convergence threshold, and the proportion of data allocated to training versus testing. Users also select the LLM for code generation, the model for literature retrieval, and the embedding model for semantic search. Finally, the user can specify a directory with curated ecological literature from which LEMMA will identify plausible parameter values and bounds. These inputs collectively define the modelling context while allowing LEMMA to operate with minimal manual intervention and substantial flexibility in model structure.

#### 2.1.3 Parameterisation

Upon initialization, the LLM estimates parameter values for each parameter in its proposed model. LEMMA uses this initial estimation to quickly run the model using TMB to identify structural or syntactical errors in the LLM-generated code. If a candidate model is successful in compiling and running, LEMMA refines these estimates using evidence from the scientific literature. Building on the success of LLM-based extraction from ecological literature (Keck et al., 2025; Spillias et al., 2024b), LEMMA implements a RAG architecture to search scientific literature (see Supplement for detailed RAG implementation). Without this initial estimation step, LEMMA risks wasting time and computational resources searching for parameter values to populate equations that may be structurally or syntactically flawed.

The RAG process works as follows: First, LEMMA prompts an LLM to create detailed semantic descriptions of each parameter, expanding beyond the basic descriptions provided by the coding LLM. For example, if the coding LLM defines a parameter as “growth rate of phytoplankton,” the RAG system might expand this to “maximum specific growth rate of marine phytoplankton in nutrient-rich conditions, measured per day.” These enhanced descriptions aim to improve the relevance of search results when querying literature databases.

To find appropriate parameter values, the RAG system employs a structured, multi-source search strategy. It searches a curated local collection of scientific papers (see Supplement), queries the Semantic Scholar database (Allen Institute for AI, 2024), and can perform general web searches through the Serper API (Serper, 2024) (although we do not employ general web-searches in the case-studies below). LEMMA combines results from all three sources to build a comprehensive understanding of each parameter’s possible values and ecological meaning.

The RAG system then uses LLMs to extract numerical values from the search results, determining not only parameter values but also their valid ranges. The prompt instructs the LLM to identify minimum, maximum, and typical values for each parameter, along with their units and citation information (see Supplement). All parameter information is stored in a structured database that includes bounds, units, and citations.

#### 2.1.4 Model Execution, Optimisation, and Error Handling

**Model Execution** Models are executed through TMB (Kristensen, 2014), which uses automatic differentiation to efficiently compute gradients for complex, non-linear optimisation problems. The dynamically generated TMB template defines the objective function internally; LEMMA treats this as a black box and does not modify its structure. We use a Control File in *R* which provides the data, starting values, and explicit box constraints for optimisation.

**Optimisation Strategy** Parameter estimation is carried out in a series of phases to improve stability and convergence, particularly in high-dimensional models. Parameters are grouped according to priorities assigned by a LLM, and only the parameters in the current group are estimated during each phase, while the others are held fixed. After all groups have been processed, a final optimisation is performed in which all parameters are free to vary.

For this proof-of-concept, all parameters are treated as estimable. Optimisation is performed using the *nlminb* algorithm, which minimises the objective function defined in the TMB (Template Model Builder) template. This optimiser uses gradient-based methods and terminates when one of its internal convergence criteria is met: either the gradient norm falls below a threshold, the parameter updates become sufficiently small, or the maximum number of iterations is reached (with a default limit of 150 iterations per phase). These stopping conditions are handled internally by *nlminb* and are not manually adjusted.

Each parameter is constrained by explicit lower and upper bounds, which are enforced at the optimiser level. These bounds, along with starting values, are derived from the literature when available; otherwise, fallback estimates provided by the LLM are used. Prior to optimisation, all starting values are clamped to lie within the admissible bounds. If a parameter’s lower bound exceeds its upper bound, the values are swapped. In cases where the bounds are equal (i.e., the interval has zero width), a small *ε* is added to expand the interval.

The optimisation is conducted under a maximum likelihood framework, leveraging TMB’s automatic differentiation to compute gradients efficiently. No regularisation terms are added by the framework; the optimisation is unconstrained apart from the explicit parameter bounds. The LLM- and literature-derived values are used solely to define starting points and bounds, and do not influence the objective function directly. Because the TMB template is generated dynamically, the framework does not impose a fixed likelihood structure, but assumes that the LLM-defined function represents a valid statistical model.

**Error Handling** Because LLMs often make trivial mistakes in their outputs, we developed an error handling system to address common issues and preserve LLM-progress, rather than discarding partial attempts. For example, on occasion, the LLM coder will attempt to create a system of equations with circular logic (“data leakage”), or will omit a key output variable with a corresponding time-series. To prevent this, we implement code validation checks to ensure that the submitted model is properly formatted and free from logical inconsistencies. For models that fail, LEMMA addresses compilation errors through automated analysis of error messages and implementation of appropriate fixes. For numerical instabilities, LEMMA employs progressive simplification of model structure while maintaining ecological relevance. Each model variant receives up to a user-specified number of fixes (five for our case-studies), with later iterations prompting for simpler model structures that can be iteratively improved. The specific prompts used for error handling are provided in Supplement.

#### 2.1.5 Model Evaluation

For each response variable *j*, we calculate a normalized mean squared error:

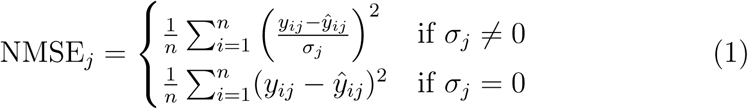

where *y_ij_* represents observed values for variable *j* at time *i*, *y*^*_ij_* represents corresponding model predictions, *σ_j_* is the unbiased standard deviation of the observed values for variable *j* (calculated with *n −* 1 denominator), and *n* is the number of observations. The final objective function value is the mean across all response variables:

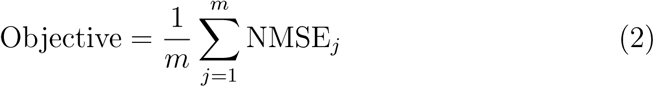

where *m* is the number of response variables. NMSE was selected as the primary objective function because it normalizes error by the standard deviation of each time-series, ensuring that all variables contribute equally to model evaluation. This avoids bias toward variables with larger magnitudes and supports balanced optimization across multivariate ecological datasets. For simplicity in this proof-of-concept, we did not weight the time-series in the objective function, however this might prove useful in future work to prioritize uncovering key dynamics.

#### 2.1.6 Evolutionary Algorithm Implementation

LEMMA maintains a population of model instances, which we refer to as ‘individuals’, where each individual represents a complete model implementation including its equations, parameters, and performance metrics. Within each generation, after mutation, individuals undergo parameter optimization using Template Model Builder to find optimal parameter values for their current model structure.

After parameter optimization, individuals are evaluated based on their prediction accuracy. Those achieving the lowest prediction errors (objective values) are selected to become parents for the next generation, while less well-performing individuals are culled and non-functioning ones (those that fail to compile or execute) are discarded.

At the beginning of each new generation, LEMMA creates new individuals in two ways: by making targeted modifications to the code in the chromosomes, model.cpp and parameters.json, of the best-performing parent individuals from the previous generation, and by creating entirely new individuals from scratch when there are not enough functioning individuals from previous generations. Both of these are done via Aider’s underlying tool handling routines that enable file modifications and creations and are determined by the underlying LLM’s response to the standard prompts, the project topic, and the current model performance (if modifying an existing model).

### 2.2 Validation Experiments

We conducted two complementary validation case studies of LEMMA. The first validation experiment aimed to see if LEMMA could recover known model equations from synthetic time-series data, whilst the second validation experiment examined real-world applicability through modelling ecological dynamics on the Great Barrier Reef. In addition to the primary experimental design (training on the full dataset), we conducted a supplementary temporal hold-out evaluation for the COTS case study using the best-performing LLM, with three replicate populations, to assess out-of-sample predictive performance.

#### 2.2.1 LLM Configuration and Experimental Design

We utilized three leading-edge LLMs to drive LEMMA’s model generation and improvement processes: GPT-5, Claude-Sonnet-4.5, and Gemini-2.5-Pro. These models were selected as they represent the current state-of-the-art in language model capabilities at the time of analysis. All models were accessed via OpenRouter, and the analysis was conducted in October 2025. To maintain consistency and reduce complexity in outcomes, we employed GPT 4.1 as a dedicated support model for all parameter estimation and RAG-related queries across all three main LLMs. This approach ensured that differences in model performance could be attributed to the core model generation capabilities rather than variations in parameter retrieval or literature search processes.

To systematically evaluate LEMMA’s performance across different LLMs and ecological systems, we implemented a balanced factorial experimental design. The design consisted of three replicates for each combination of the three LLMs and the two ecological case studies (NPZ and COTS). This yielded a target of 18 populations (3 LLMs *×* 2 case studies *×* 3 replicates). Each population was initialized with four individuals and allowed to evolve for a target of ten generations, unless convergence was reached earlier (defined as achieving an objective value improvement of less than 0.01 for three consecutive generations).

Because we use frontier, proprietary LLMs, LEMMA can be run on a conventional laptop and LLMs can be accessed via API calls, which effectively outsources the heavy computation needed for LLMs at a financial cost. During development we succeeded in running LEMMA with open-source LLMs on local hardware (specifically GPT-OSS:120b on a a single H100 GPU), however because it was not connected to the internet, equivalent parameterisation, including Semantic Scholar database searches, was not possible and we therefore do not report those results here.

#### 2.2.2 Retrieving Model Equations – NPZ Case Study

We conducted a controlled experiment using synthetic time-series data generated by a well-established nutrient-phytoplankton-zooplankton (NPZ) model from Edwards and Brindley (1999), whose dynamics are well-studied (Boschetti, 2008, 2010). The complete system of equations is presented in S3.5. This validation tested our framework’s ability to rediscover established ecological relationships from synthetic data where the underlying equations of a system are known, providing a rigorous assessment of the system’s equation-learning capabilities.

In addition to monitoring the convergence of LEMMA’s modelled time-series towards the provided time-series data, we evaluated the framework’s ability to recover nine key ecological characteristics from the original model. These characteristics were organized by equation: three components in the nutrient equation (dN/dt: uptake, recycling, mixing), four components in the phytoplankton equation (dP/dt: growth, grazing loss, mortality, mixing), and two components in the zooplankton equation (dZ/dt: growth, mortality). For each individual, we used GPT-5-mini to evaluate each model component using a 4-level ordinal scoring system designed to be interpretable and verifiable by ecological experts:

- TRUTH_MATCH: The mathematical structure is equivalent to the ground truth NPZ model
- ALTERNATE: The implementation matches a recognized alternate formulation from the ecological literature (e.g., different functional response curves from (Franks, 2002))
- SIMILAR_NOT_LISTED: The implementation plays the same ecological role but uses a form not represented in the ground truth or literature catalog
- NOT_PRESENT_OR_INCORRECT: The ecological component is missing or cannot be identified

The complete evaluation prompt with detailed scoring criteria and catalog of alternate formulations is provided in Supplement. This additional evaluation allowed us to understand the extent to which LLMs were relying on established ecological understanding or generating equations unsupported by ecological theory.

To complement time-series convergence and ecological scoring, we compared the structural forms of individual process components between the ground-truth NPZ model and the best-performing LEMMA-derived model. For this comparison, each flux term was plotted against its primary driver while holding other state variables constant at representative initial values. Optional modifiers present in the compiled model (e.g., temperature and light limitation, quadratic zooplankton mortality, detrital recycling) were disabled to isolate differences in core equation structure. Parameter values were aligned between models so that observed discrepancies reflect functional form rather than parameterization.

#### 2.2.3 COTS Case Study

The Crown-of-Thorns starfish (COTS) case study examined real-world applicability through modelling populations of COTS and their prey, coral, on the Great Barrier Reef. This case study also made two external forcing time-series available to LEMMA, sea-surface temperature and COTS larval immigration quantities. We tested the three main LLMs (GPT-5, Claude-Sonnet-4.5, and Gemini-2.5-Pro) and evaluated LEMMA’s ability to match the model created by a human expert in the same context.

The COTS model that we used as a human-derived benchmark for evaluating LEMMA’s outputs was originally developed to specifically evaluate management interventions (Morello et al., 2014). Subsequent model versions and variations have yielded insight into management under environmental perturbations (Rogers and Plagányi, 2022; Condie et al., 2021), derivation of management thresholds (Plagányi et al., 2020; Rogers et al., 2024) and their dynamic implementation (Rogers et al., 2023). Each application differed depending on the objectives and data availability, and how the models were resolved, which required human determinations as to what was included, how it was included, and how it linked with other system aspects where necessary. Simply put, human experts were required to link management objectives and available data to resolve the necessary system aspects for informing specific management actions.

Here we test LEMMA’s ability to develop a model that captures the dynamics of corals and COTS during a COTS outbreak. This not only required LEMMA to link the ecological dynamics but also interpret the life history characteristics of COTS to explain the observed data. Due to time and cost constraints we only performed limited tests using each LLM, where we initialized populations of four individuals for ten generations each. We calibrated these population parameters by balancing the cost of running an individual population against the rate of convergence that we found in our initial tests of the system.

We tracked several key performance metrics for each population:

- Runtime performance: Total runtime and per-generation computation time
- Error resolution: Number of iterations required to achieve successful model implementation in each generation
- Model stability: Proportion of successful, culled (underperforming), and numerically unstable models per generation

We also analyzed the evolutionary trajectories of successful models by tracking their lineage from initial to final states, documenting the frequency and magnitude of improvements across generations. This included measuring the number of generations required to reach best performance and the proportion of attempts that resulted in improved models.

We performed a temporal hold-out evaluation by partitioning the time series into a training period (pre-1997, approximately 70%) used for parameter estimation and a test period (1997-2005; 30%) used once for out-of-sample assessment. For each ecosystem component (COTS abundance, fast-growing coral cover, and slow-growing coral cover), we calculated root mean square error (RMSE), mean absolute error (MAE), and R² values to quantify prediction accuracy. By comparing these metrics against those of the human-developed reference model, we could assess whether our automated approach could match expert-level performance in a real-world ecological application.

## 3 Results

### 3.1 Retrieving Model Equations – NPZ Case Study

LEMMA achieved strong success in recovering the quantitative fit of the NPZ test model across all tested LLMs within ten generations. GPT-5 achieved the most accurate results, with three populations reaching objective values of 0.004, 0.007, and 0.023 (lower is better), indicating rapid convergence and high predictive precision (with two populations converging below 0.01 in 3 and 7 generations). Gemini-2.5-Pro showed mixed outcomes, with its best population at 0.036, but others at 0.377 and 1.881. Claude-Sonnet-4.5 delivered intermediate performance, with objective values ranging from 0.153 to 0.787. Overall, GPT-5 populations dominated in terms of convergence and error minimization, while Gemini and Claude provided reasonable approximations but with greater variability (Figure 3).

**Figure 3:**
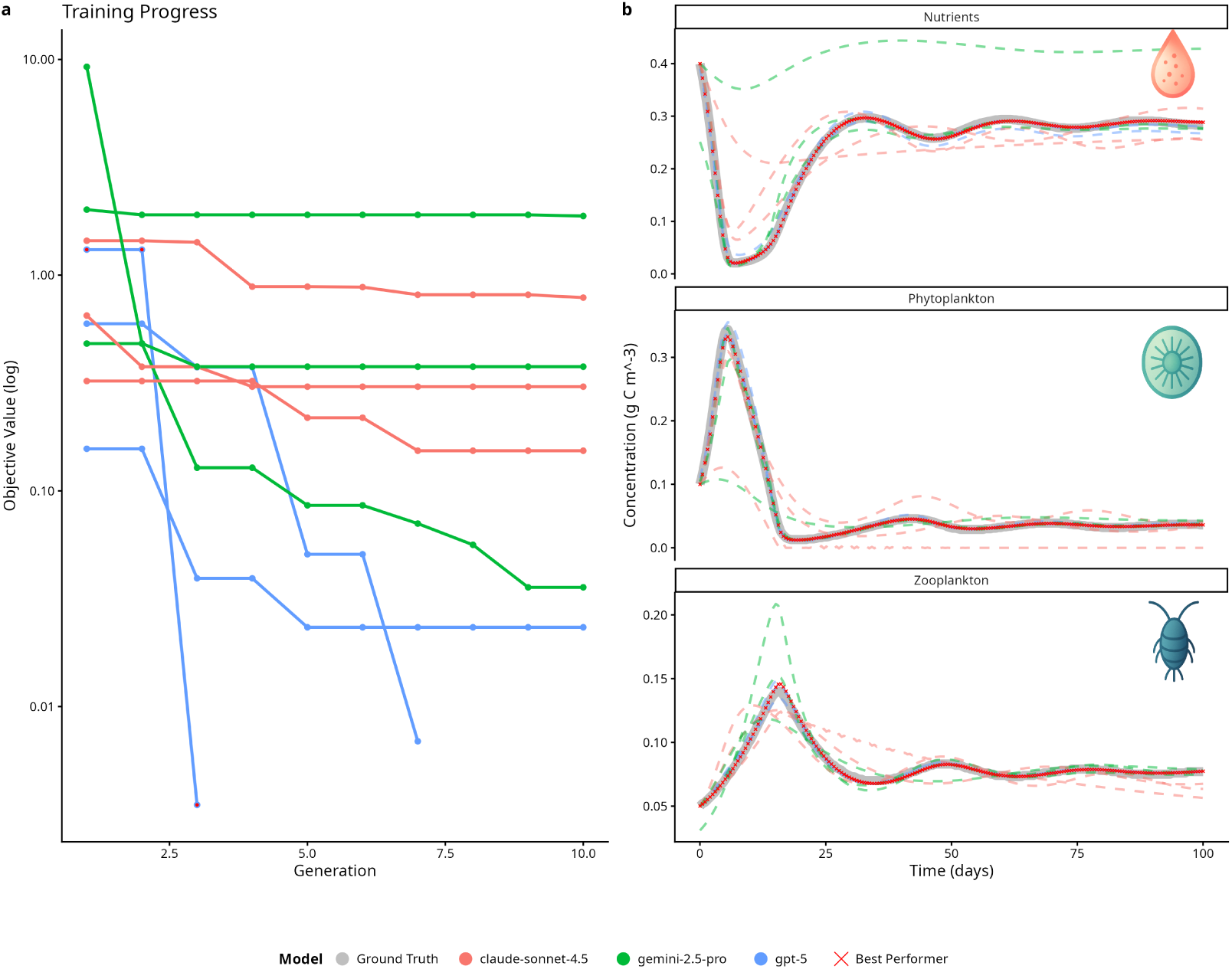
Performance and ecological dynamics of NPZ model retrieval. (a) Training progress showing objective value trajectories across generations (logscale) for all populations driven by GPT-5, Claude-Sonnet-4.5, and Gemini-2.5-Pro. GPT-5 achieved the lowest objective value (0.004) within three generations. (b) Time-series comparison of nutrient, phytoplankton, and zooplankton concentrations (g C m^-3^) between ground-truth (black lines) and predictions from the best model in each population by objective value (crosses). The model successfully reproduced bloom timing, nutrient drawdown, and trophic phase relationships, indicating strong alignment with ecological mechanisms.

Recovery of NPZ mechanisms varied substantially across LLMs. No model individual LEMMA model perfectly reconstructed all processes, but every mechanism was recovered at least once (Figure 4). GPT-5 achieved the highest overall proportion of well-supported formulations (alternate from Franks (2002) or truth match), with well-supported mechanisms accounting for 80.8% overall versus 19.2% not supported. For the best-performing individual (GPT-5), only phytoplankton mixing was unsupported (not a truth match or alternate). With the exception of both mixing processes, all of the LLMs tended to use supported formulations. Quantitatively, the fluxes of most of the formulations used by the best-performing model were exact or very close matches to those of the truth model. However, the best-perfomer also included a detritus pathway that was not present in the ground-truth (Figure 5). These patterns indicate that while overall ecological fidelity was high, certain processes, like phytoplankton and nutrient mixing were often difficult (but not impossible) for LLMs to reconstruct accurately and that quantitative matches were substantiated by ecological intention.

**Figure 4:**
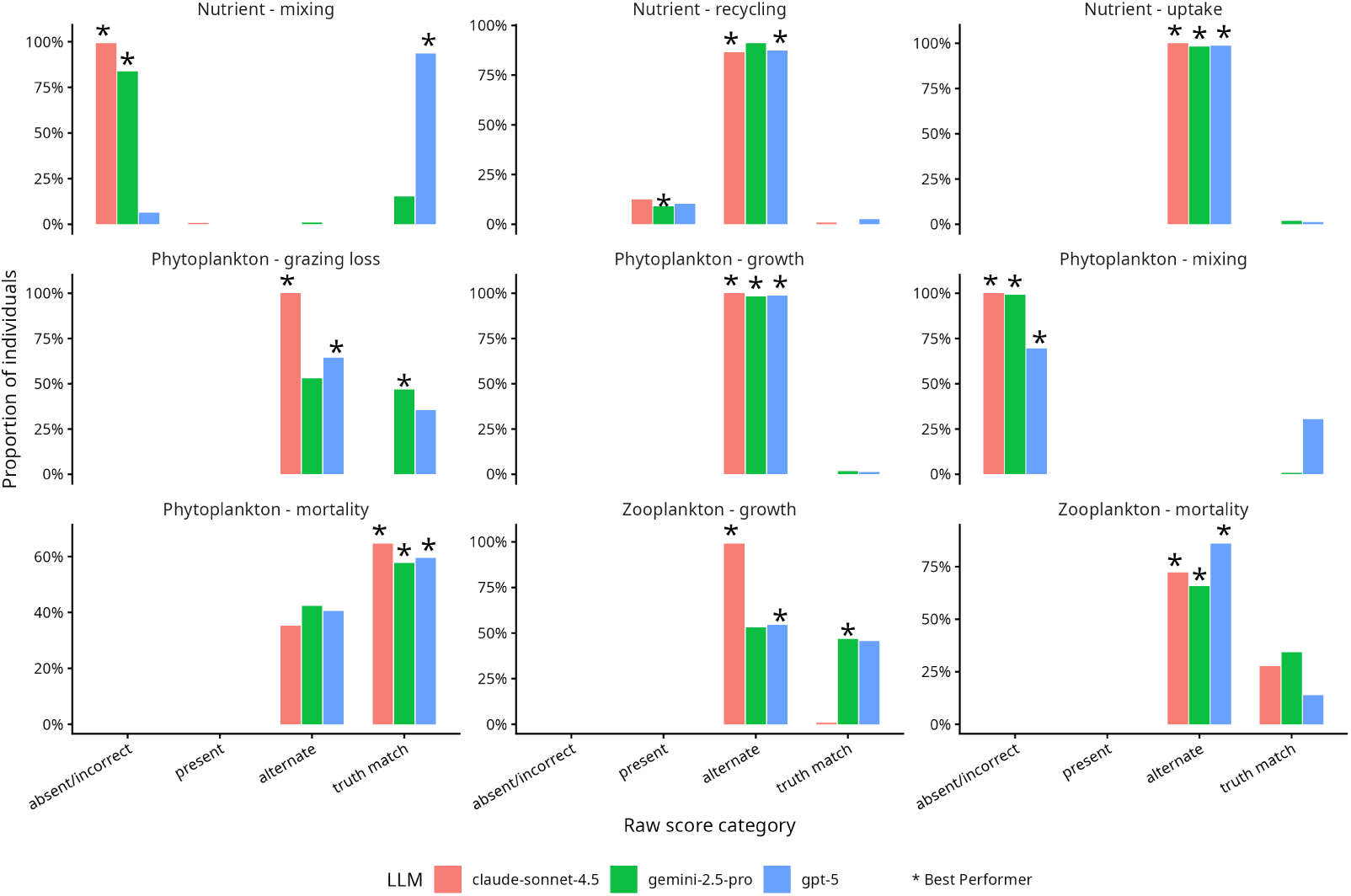
Mechanism-level recovery of NPZ equations across LLM families. Each panel shows the distribution of raw scores for a specific ecological mechanism as described in Section 2.2.2

**Figure 5:**
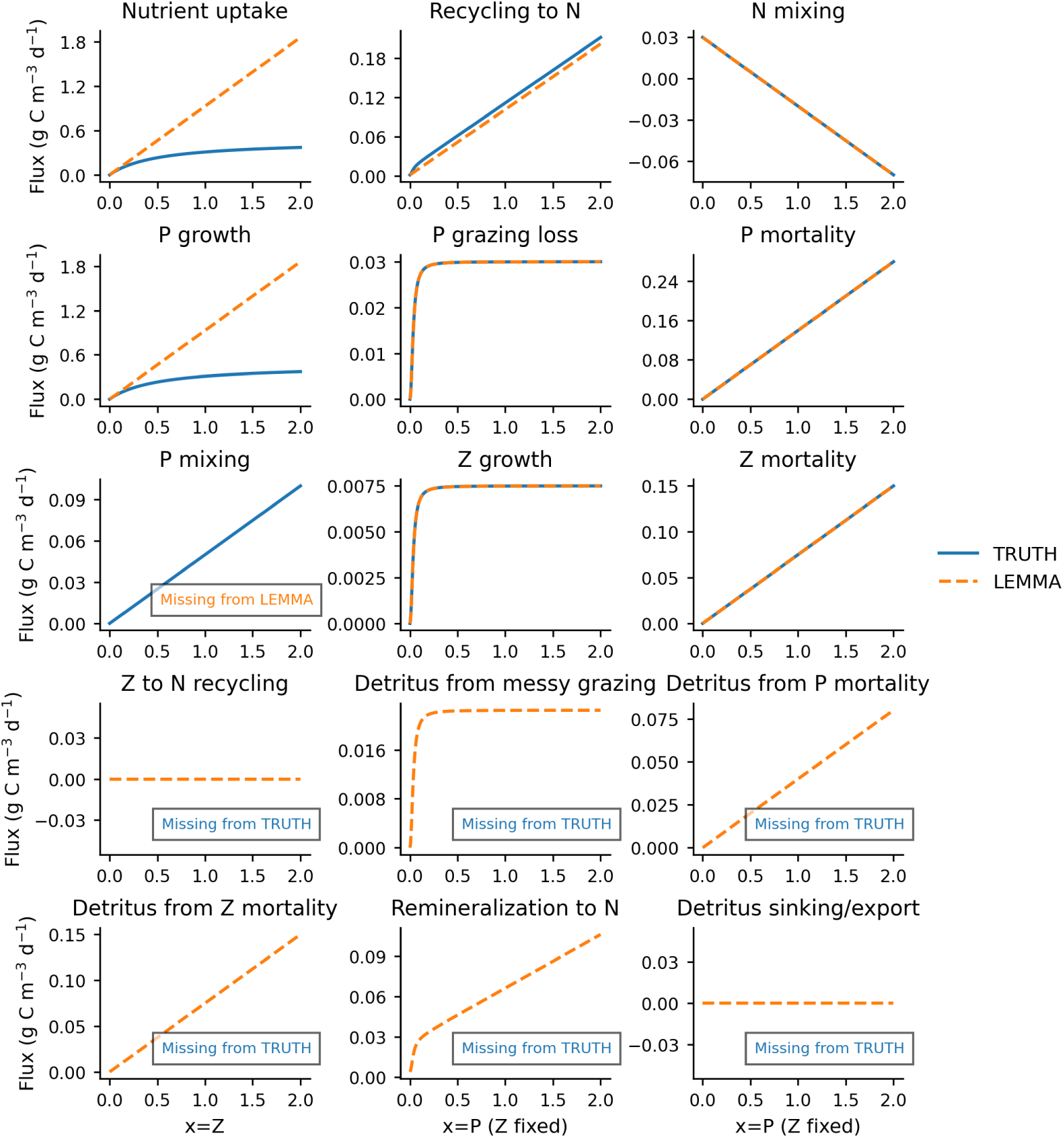
Comparison of process-level fluxes between the ground-truth NPZ model and best-performing LEMMA-derived model. Each panel shows the functional form of a single flux term plotted against its primary driver, with other state variables held constant at representative initial values. Optional modifiers in the compiled model (temperature, light, quadratic zooplankton mortality, detrital recycling) are disabled to isolate core equation differences. Parameter values are aligned so that curve differences reflect structural form rather than parameterization. Processes absent in one model are indicated as missing.

In addition to the presence of the ground-truth processes, we also saw that the best-performing LEMMA model successfully reproduced most ecological dynamics of the NPZ system according to the specific fluxes, although it also included additional processes that were not present in the ground truth model (Figure 5). Uptake, phytoplankton growth, and zooplankton mortality exhibited near-perfect alignment with the ground-truth model, indicating that core trophic interactions were accurately reconstructed. Grazing loss and recycling processes were also well captured, though minor deviations were observed in recycling to nitrogen at higher flux values. As reported above, phytoplankton mixing was absent in the LEMMA-derived model. The best-performing model also included a detritus pathway which was not present in the original ground-truth model.

### 3.2 COTS Case Study

For all tested LLMs, LEMMA was able to generate ecosystem models with prediction accuracy approaching the quantitative fit of the expert-developed model (objective value NMSE: 0.2312). After ten generations, objective values across LLMs (based on three independent runs per model) were as follows: GPT-5 achieved the best performance, with objective values ranging from 0.31-0.45 (lower is better), followed by Sonnet-4.5 (0.33-0.44) and Gemini-2.5-Pro (0.47-0.69). Component-specific analysis revealed varying levels of prediction accuracy, with models showing strongest performance in predicting fast-growing coral cover and slow-growing coral cover, while maintaining reasonable accuracy for the more volatile COTS abundance patterns (Figure 6). GPT-5 was able to correctly model the outbreaks of COTS, as requested in the initial prompt, while Gemini-2.5-Pro’s best model was out of phase and Sonnet-4.5’s identified the first, but not the second.

**Figure 6:**
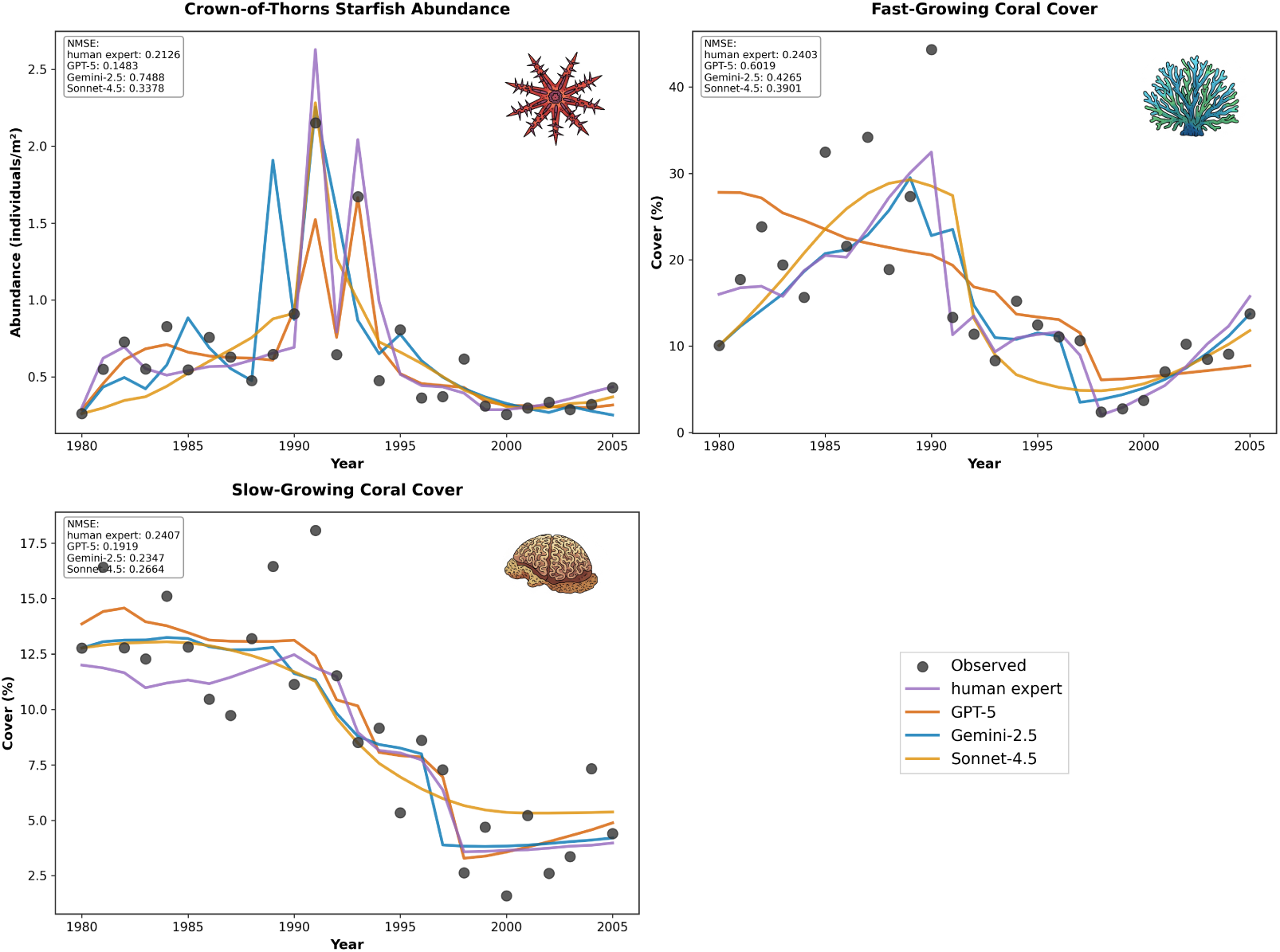
Comparison of model predictions across ecosystem components. The plots show observed versus predicted values for COTS abundance, fast-growing coral cover, and slow-growing coral cover, demonstrating the models’ ability to capture key ecological patterns and relationships. Objective values (obj) shown in the legend represent the normalised mean squared error for all three variables, where lower values indicate better model performance.

The best-performing models generated by GPT-5, Claude-Sonnet-4.5, and Gemini-2.5-Pro exhibited differences in structural design and ecological mechanisms, whilst also sharing several elements. GPT-5 aligned most closely with the demographic logic of the human reference model, while Gemini and Claude adopted more simplified, outbreak-oriented frameworks. Across all LLMs, certain core features remained consistent: density-dependent population regulation in crown-of-thorns starfish (COTS), resource limitation driven by coral availability, and differential effects on fastversus slowgrowing coral species. These commonalities occurred alongside substantial variation in life-history representation, recruitment formulation, immigration handling, mortality drivers, predation dynamics, and thermal stress responses (see Supplement).

Among the LLM-generated models, GPT-5 most closely resembled the human reference model in population structure. GPT-5 modelled crown-of-thorns starfish (COTS) using two stages, juveniles and adults, while the human model applied a more detailed age-structured approach with three classes and age-specific mortality rates. In contrast, Gemini simplified the population to a single undifferentiated compartment, and Claude used a single adult compartment supplemented by episodic recruitment pulses. For recruitment, GPT-5 incorporated a Beverton-Holt-like taper on adult abundance, modified by environmental factors and lagged immigration, which paralleled the continuous Beverton-Holt density dependence used in the human model. Gemini, on the other hand, linked recruitment to coral consumption and introduced Allee effects and temperature windows without an explicit stock-recruitment curve. Claude adopted a threshold-based pulse mechanism rather than a continuous function. GPT-5 handled immigration by integrating with recruitment and applied a temporal lag, echoing the demographic embedding seen in the human model, which introduced immigration during age-0 formation with lognormal variability. Gemini instead added immigrants directly to the adult compartment, while Claude combined immigration with favorability indices and growth multipliers, creating a compound influence on population dynamics. GPT-5 and Gemini both applied baseline mortality augmented by density-dependent feedbacks, which partially reflected the human model’s approach of age-specific mortality tied explicitly to prey availability. Claude introduced a starvation multiplier instead that amplified mortality when coral cover was low. For predation, GPT-5 blended Holling Type II and Type III responses, producing low-prey refuges, whereas the human model used a sigmoid saturation curve with explicit prey-switching rules. Gemini employed a standard Holling Type II functional response, and Claude incorporated prey preference weighting rather than explicit switching. For coral growth and thermal stress, GPT-5 penalized coral growth exponentially above a thermal threshold and included a heat-loss term, which partially mirrored the human model’s use of Gaussian performance curves for growth combined with logistic bleaching mortality. Gemini restricted thermal effects to logistic bleaching only, while Claude applied linear mortality beyond stress thresholds without Gaussian modulation. Finally, temperature effects on COTS recruitment were absent from the human model but present in all LLM implementations. GPT-5 and Gemini used Gaussian temperature functions to modulate recruitment, whereas Claude embedded temperature influences within outbreak pulses and adult growth factors.

### 3.3 Evolutionary Behaviour

There were notable differences in the behaviour and speed of convergence across all LLMs tested (Supplemental Figure 1). The Gemini-2.5-Pro models were the fastest overall, completing ten generations in approximately 36-50 minutes, with a mean time per generation of just over four minutes. GPT-5 models showed intermediate performance, averaging about 12 minutes per generation.

Patterns of improvement also differed markedly. Sonnet-4.5 and Gemini-2.5-Pro generally produced well-functioning models early but struggled to achieve consistent gains across generations, with mean improvement rates close to zero and frequent plateaus. GPT-5 demonstrated more stable progress, including the best-performing population overall (objective value 0.004), though most GPT-5 runs also exhibited diminishing returns after initial improvements.

#### 3.3.1 Temporal Hold-Out

Because GPT-5 demonstrated the strongest overall performance, we used it as the base LLM to test whether the LEMMA framework could create models capable of predicting out-of-sample datapoints. After ten generations, we found that the best-performing model (objective value: 0.29) had a reasonable degree of out-of-sample prediction performance on the withheld test dataset, with particularly strong predictive power for fast-growing coral cover (R² = 0.81, RMSE = 2.22, MAE = 1.92). Interestingly, this was a better objective value than the previous experiments where LEMMA had access to the entire time-series.

For slow-growing coral cover, the model achieved moderate predictive accuracy (R² = 0.17, RMSE = 1.6, MAE = 1.3), effectively capturing the general declining trend while showing some deviation in precise values. COTS population predictions demonstrated strong accuracy (R² = 0.75, RMSE = 0.07, MAE = 0.05), successfully capturing the timing and magnitude of major outbreaks.

Figure 7 illustrates these prediction capabilities, showing both training period performance (pre-1997) and out-of-sample predictions (1997-2005). The model’s ability to maintain consistent error metrics (RMSE and MAE) while capturing both rapid population dynamics and slower coral cover changes suggests it successfully identified fundamental ecological relationships governing this system.

**Figure 7:**
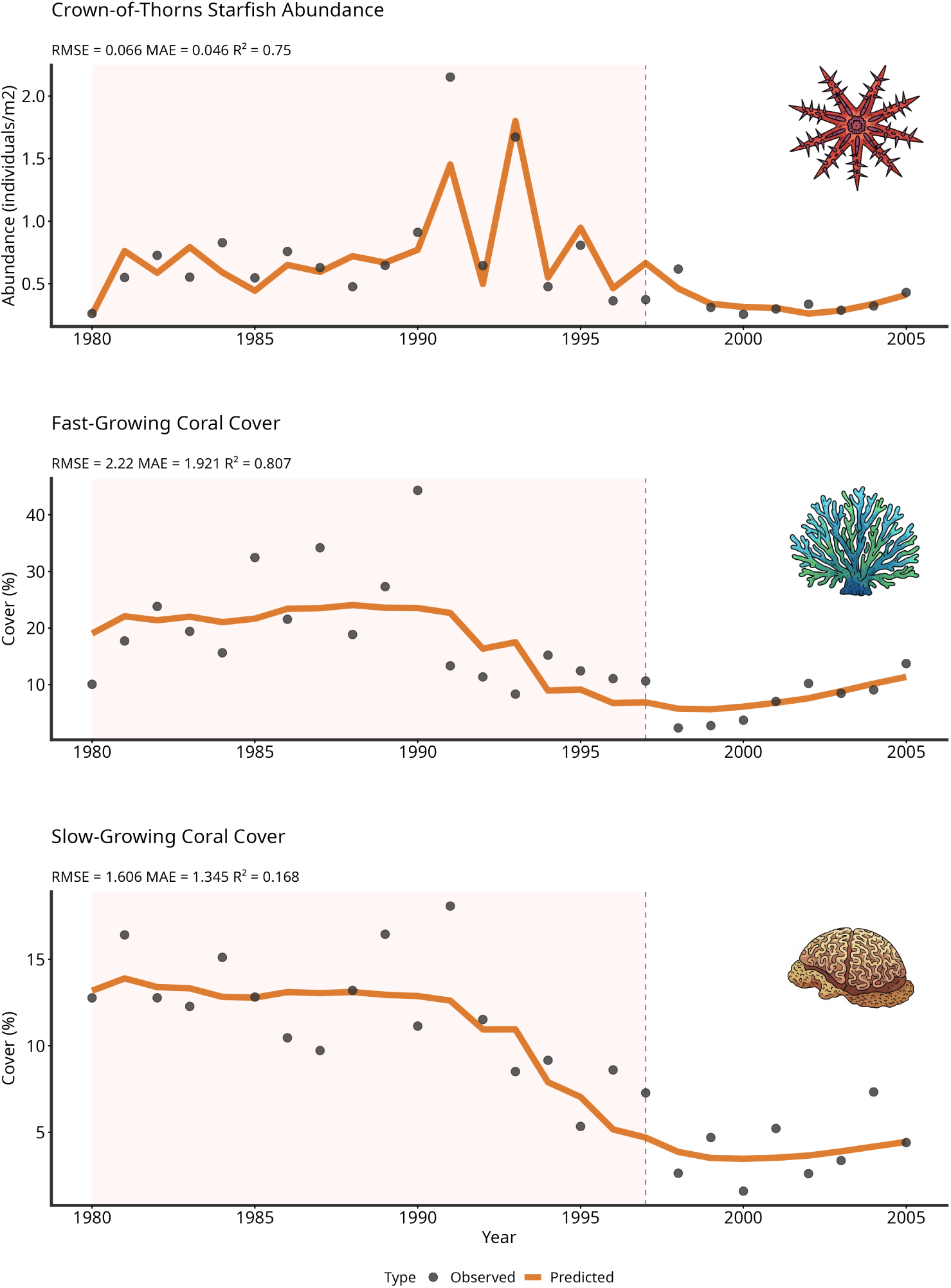
Temporal hold-out evaluation of LEMMA driven by the best-performing LLM (GPT-5) showing predictions against observed data. LEMMA was provided with 70% of the time-series data (pink shaded region) and developed a model that was then evaluated on the full time-series, including the remaining unseen 30% of the time-series data (white region). Lines represent model predictions, dots show observed data.

**Figure 8:**
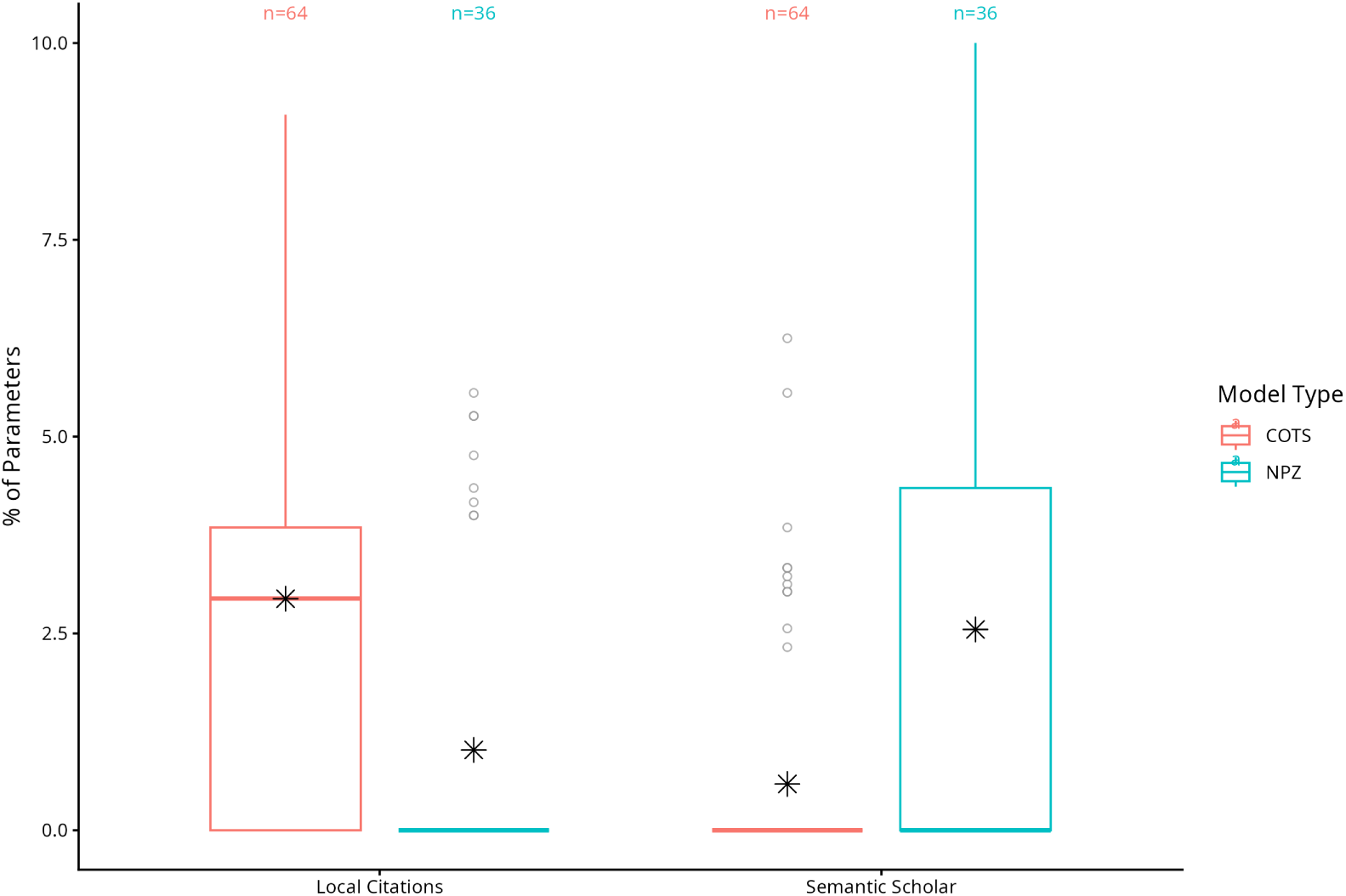
Comparison of citation integration between NPZ and COTS case studies. Boxplots illustrate the proportion of model parameters with citations (Semantic Scholar or local document store). NPZ models were more reliant on Semantic Scholar because the local document store was curated specifically for the COTS case study. Asterisks represent the best performer (lowest nRMSE objective value) for each model type.

### 3.4 Parameter Retrieval

We assessed how frequently LEMMA linked model parameters to external evidence sources, including Semantic Scholar searches and a locally curated document store. Across all generated models (21 populations, 2,619 parameters), citation integration was modest: only 3.1% of parameters were linked to explicit citations, 1.2% to Semantic Scholar matches, and 2.0% to document store references. The majority of parameters lacked any external linkage. Differences emerged between case studies. NPZ-based populations showed slightly higher citation rates (3.7%) and semantic matches (2.7%) than COTS-focused populations (2.8% and 0.59%, respectively). Document store usage was notably higher for COTS models (2.4%) compared to NPZ (1.1%). This disparity likely reflects the fact that the document store was curated specifically for the COTS case study, whereas no curation was performed for NPZ, limiting retrieval opportunities. At the individual model level, citation coverage rarely exceeded 10% of parameters, and many individuals contained no cited parameters. Where citations were present, they typically supported temperature optima, growth coefficients, and bleaching thresholds, parameters with well-documented ecological relevance. These results indicate that while LEMMA can incorporate literature-based evidence, this capability was underutilized in practice. Enhancing citation integration may require prioritizing literature-grounded parameter selection and expanding curated resources for all ecosystem contexts.

## 4 Discussion

Our complementary validation studies demonstrate the viability of AI-driven automation in ecological model development and calibration. The NPZ case study revealed LEMMA’s ability to achieve very high quantitative fidelity whilst also recovering known ecological relationships from synthetic data. While LEMMA did not perfectly reconstruct the original equations after ten generations, it incorporated all ecological mechanisms except phytoplankton mixing; with the remaining components being either identical to the NPZ reference model or accepted alternatives from Franks (2002). Whilst we cannot rule out the possibility that this success is a result of this NPZ model (and those like it) being well-studied and common in the literature - and therefore likely in the training data of these LLMs - this outcome nonetheless demonstrates the ability of LLMs to retrieve and adapt ecologically meaningful models. Importantly, the best-performing LEMMA-derived models did not simply reproduce the NPZ equations verbatim. They included alternate formulations and additional mechanisms such as detritus pathways, which were not present in the original model. This suggests that the LLMs are capable of generalizing from known ecological principles rather than merely memorizing specific models. We interpret this as evidence that LLMs can transfer mechanistic understanding across contexts, a capability that may prove valuable for accelerating ecological model development in practice.

In the COTS case study, LEMMA-generated models achieved predictive performance that was broadly similar to that of the human expert model, although with various structural differences. While the human model implemented an age-structured COTS population with three age classes, only LEMMA with GPT5 incorporated a size-structured COTS population (with two classes). Similarly, the human model featured an explicit prey-switching function for COTS predation preference between coral types, whereas LEMMA models used Holling Type II/III responses, preference weighting, and coral-dependent modifiers, but did not replicate the explicit switching function. This is consistent with the coexistence of multiple numerically valid representations of the same ecological system where each offers different insights into the underlying mechanisms (Patterson et al., 2001).

This flexibility also opens the door to hybrid approaches that combine the strengths of both expert-driven and AI-generated models. By explicitly prompting LEMMA to include particular ecological processes, such as age structure, prey-switching behavior, or nutrient mixing, researchers can guide model structure to reflect known system dynamics or stakeholder priorities. This capability enhances the utility of AI-generated models for hypothesis testing, scenario exploration, and applied decision-making, particularly in contexts where certain mechanisms are known to be ecologically or socially important.

Whilst it is difficult to be certain exactly what is in any given LLM’s training data, we can safely assume that their embedded ‘knowledge’ will be roughly correlate to that which is commonly found on the internet Brown et al. (2020). Therefore, it is likely that LEMMA will work best in well-studied systems where data and analysis is plentiful and may struggle in systems that are poorly documented and understudied. However, it is worth distinguishing between two classes of understudied systems. The first would be systems that are understudied but which are highly analogous to well-studied systems (e.g., coral reefs, seagrass beds or mangroves in under-studied regions of the Global South). In such cases, LEMMA will likely be very useful at adapting existing modelling frameworks to these new places. The second class would be systems that are understudied and also lacking in well-studied analogues (e.g., deep-sea systems). In these cases, LEMMA will have less to draw from and may be less accurate, although it should not be dismissed as a potential way of generating hypotheses that can be tested via empirical study.

### 4.1 Contrasting Approaches to AI in Ecological Modelling

Recent advances in AI have demonstrated remarkable capabilities in ecological time-series prediction. Studies using transformer architectures and diffusion models, including multimodal approaches like LITE (Li et al., 2024), have shown high accuracy in direct forecasting of environmental variables (Morales-García et al., 2024; Gandhi et al., 2024). While these methods effectively handle challenges like missing data and distribution shifts, they treat the system as a black box, learning patterns directly from time-series data without explicitly modelling underlying mechanisms. While our study did not directly compare our approach with black-box methods, our NPZ validation study suggests a key advantage of our approach: the ability to provide insights into fundamental ecological processes like nutrient cycling or predator-prey dynamics through explicit model discovery. This interpretability is a theoretical advantage over black-box approaches, though comparative studies would be needed to fully evaluate the relative strengths of each approach in specific ecological contexts.

Our evolutionary approach fundamentally differs by using AI to generate actual ecological models rather than make direct predictions. Instead of training neural networks to forecast future values, LEMMA evolves interpretable models with meaningful parameters that capture real biological and physical processes. This distinction is crucial for several reasons. First, our generated models provide scientific insight into system behavior, revealing mechanisms and relationships that direct prediction approaches typically cannot without careful oversight (Adams et al., 2017). Second, the models maintain biological plausibility through explicit parameter constraints and mechanistic formulations, ensuring their utility for management applications. Third, because they capture fundamental processes rather than just patterns, these models can potentially be transferred to new scenarios and used to explore management interventions. The relationship between our approach and direct prediction methods is nuanced. Recent time-series prediction approaches using transformer architectures have achieved impressive accuracy, with mean squared errors as low as 0.001-0.04 for normalized pre-dictions (Morales-García et al., 2024) and root mean squared errors reduced by up to 52% compared to traditional methods (Gandhi et al., 2024). While our evolved models may not always match these pure prediction accuracies, they offer advantages in interpretability, scientific insight and potential for strategic and tactical applications.

Importantly, these approaches need not be viewed as mutually exclusive. Comparing mechanistic models with black-box predictions can be particularly insightful, especially when the two approaches diverge. For instance, under novel conditions like future climate scenarios, differences in predictions could highlight processes that are not well-captured by mechanistic models or reveal patterns that black-box approaches detect but cannot explain. When predictive approaches outperform mechanistic models, this divergence can guide researchers toward missing processes or relationships that should be incorporated into mechanistic understanding. Our framework demonstrates that it’s possible to achieve both reasonable predictive accuracy and meaningful ecological interpretability, with each approach offering complementary strengths.

### 4.2 Limitations and Future Directions

Despite promising results, several limitations warrant consideration. The observed variation in convergence rates across populations suggests that initial conditions significantly influence model evolution trajectories and LEMMA may get trapped in local minima. Whilst it is possible that this actually mirrors the conventional approach that a human modeller would take, exploring other forms of ‘mutation’ to increase individual diversity may lead to more consistent outcomes.

Another important limitation in our current implementation is the treatment of all model parameters as estimable quantities in the optimization process, even when well-established values exist in the literature. While our RAG system successfully retrieves literature-based values and ranges, these are used as initial estimates and bounds rather than fixed quantities. This design choice reflects the reality that literature values may not always be appropriate for the specific ecological context being modeled. Estimating parameters within literature-derived bounds allows the model to adapt to system-specific deviations, offering greater flexibility and ecological realism. Future versions of the framework could distinguish between parameters that truly require estimation and those that could be fixed, reducing the parameter space and improving computational efficiency. This would not only reduce the parameter space for optimization but also better incorporate established ecological knowledge into the modelling process and make the process less resource-intensive when doing calculations.

Parameter provenance remains a critical limitation in our current implementation. Although we restricted retrieval to curated local literature and Semantic Scholar abstracts to ensure reliance on peer-reviewed sources, only about 3% of parameters were linked to explicit citations. This gap is partly expected because many parameters are scaling constants or mathematical utilities rarely documented in the literature, and full-text articles are often locked behind paywalls. When literature values were unavailable, LEMMA defaulted to LLM-derived estimates, which, while often plausible, lack explicit citation. These limitations underscore the importance of human oversight: experts should supplement LEMMA’s outputs to ensure that key biological parameters are grounded in empirical evidence while computational parameters remain transparent. Notably, LLMs have demonstrated strong potential for extracting ecological data at scale (Gougherty and Clipp, 2024; Keck et al., 2025). Future iterations could leverage these capabilities by enabling dynamic access to curated databases, applying confidence scoring, and cross-validating retrieved values against multiple sources. Such enhancements would strengthen parameter provenance, reduce reliance on uncited estimates, and improve transparency and scientific rigor.

Unlike traditional machine-learning approaches which use regularisation and early-stopping to avoid overfitting. LEMMA employs two nested optimisation processes, both of which may need mechanisms to address overfitting. For parameter estimation within TMB, literature-informed bounds act as weak regularisation, but no additional penalties are applied. For structural evolution, we limit changes to one modification per generation and cap the number of generations. Nonetheless, explicit complexity penalties or information-theoretic criteria (e.g., AIC/BIC) could further reduce overfitting risk in future iterations.

There are numerous future avenues for validating and extending this framework. First, there are several meta-parameters that likely control the success and speed of convergence of the framework (LLM-choice, LLM temperature setting, number of individuals per generation, prompt construction, etc.). Systematic testing across these choices may reveal optimal configurations for convergence. In particular, the comparative analysis of different AI configurations (as detailed in section S5.1 and Supplemental Figure 1) reveals trade-offs between model choice and rate of improvement. While GPT-5 demonstrated the most stable improvement and achieved the best overall performance, Sonnet-4.5 and Gemini-2.5-Pro often produced strong initial models but did not consistently improve across generations. Future work could explore hybrid approaches that leverage the strengths of different LLMs at various stages of model development, or employ different models consecutively over multiple generations. Further, ongoing testing of new LLMs as they are released may yield considerable gains in efficiency and cost-saving. Second, we have tested a relatively simple ecosystem model with three dependent variable time-series and two forcing variable time-series. Simple systems like these will be limited in real-world utility, and therefore testing on more complex systems with tens or hundreds of time-series will be needed. Incorporating spatial components may also be possible and will greatly improve the utility of this framework. Third, accessing relevant scientific information for the parameter RAG search is limited by the user’s ability to either curate a local database of relevant materials, or access scientific papers online. Fourth, future work could explore alternative modelling paradigms to the forward simulation method explored here, such as state-space formulations (Auger-Méthé et al., 2021), gradient matching (Ellner et al., 2002), or one-step-ahead prediction objectives (Munch et al., 2023). Finally, our evaluation approach relies mainly on error-based metrics, which may not fully reflect ecological plausibility. Recent work on LLM-based evaluation policy extraction (Cheng et al., 2025) offers a promising direction for incorporating expert-informed, interpretable criteria into automated workflows. Future versions of our framework could integrate such methods to improve ecological relevance in model selection.

### 4.3 Implications for Ecosystem-Based Management

Acknowledging these limitations and opportunities for future development, LEMMA has clear potential to support Ecosystem-Based Management (EBM) (Pikitch et al., 2004), provided key practical considerations are addressed. In EBM, mechanistic understanding is as critical as predictive accuracy, enabling stakeholders to evaluate credibility and trace intervention pathways (Persson et al., 2014). By producing explicit equations and parameters, LEMMA promotes mechanistic reasoning and transparent scrutiny of assumptions. However, integrating LEMMA into management workflows introduces both opportunities and risks.

One major opportunity lies in accelerating model development. Traditional workflows often rely on repurposing legacy code, which can embed hidden assumptions poorly suited to new contexts. LEMMA could enable rapid construction of models from first principles, making assumptions explicit and auditable. This capability would also better align model development timelines with tactical decision-making needs. Reduced time and effort would also allow modellers to replicate the model-building process without bias from prior implementations, which is an important but rarely feasible aspect of scientific reproducibility. By lowering technical barriers, LEMMA might also empower non-modellers and stakeholders to create models informed by local or traditional knowledge, deepening engagement and contextual understanding. Further, LEMMA’s tendency to propose ecological concepts absent from human-derived models highlights its potential for hypothesis generation and discovery of novel mechanisms. These ideas could inform new research directions and targeted data collection, strengthening the adaptive cycle of learning and management.

However, these benefits come with challenges. Non-experts may lack the skills to evaluate outputs, verify provenance, or diagnose failure modes. And while experienced modellers would not share these challenges, using LEMMA might generate a proliferation of plausible models with uncertain ecological or policy relevance. For example, some of the temperature-related components LEMMA introduced in the COTS case study lack empirical support. And while some concepts may prove unfounded and incorrect, others may warrant empirical investigation and targeted data collection. Developing safe and efficient ways to screen these outputs for validity will be important to avoid simply shifting the capacity bottleneck from model development to model analysis and selection.

Maintaining expert oversight is essential. Automated scientific discovery emphasizes coupling AI’s computational power with human judgment (Kramer et al., 2023; Spillias et al., 2024a), and LEMMA’s interpretable outputs should enable experts to screen models that include mistaken components or miss key known-processes, thereby efficiently narrowing a large set of candidate models to a robust and diverse subset. These models can then be combined into ensembles to quantify structural uncertainty (Baker et al., 2017; Gårdmark et al., 2013; Vollert et al., 2024). And whilst we use an objective function in this study that prioritises forecast accuracy, LEMMA could be customized to optimise a range of different skill metrics (potentially simultaneously) in order to mitigate the “forecast trap” described by (Boettiger, 2022), where decisions optimized for forecast accuracy may underperform in utility space.

To ensure safe and effective integration of LEMMA into ecosystem-based fisheries management, we recommend a set of best practices and propose that LEMMA would work best embedded within a human-driven workflow (Figure 9).

1. **Stakeholder Engagement** Begin with stakeholder engagement to inform problem framing and define the ecological questions LEMMA will address. Early involvement ensures models are relevant to management needs and incorporate local knowledge.
2. **Expert Review** All AI-generated models should be reviewed by domain experts to validate ecological plausibility and ensure alignment with management objectives.
3. **Complementary Use** Use LEMMA as a complementary tool that supports rather than replaces traditional modelling workflows, especially for rapid prototyping or exploring alternative hypotheses.
4. **Model Transparency** Maintain transparency through clear documentation of equations, parameters, and assumptions to ensure traceability and reproducibility.
5. **Parameter Assessment** Critically assess parameter values sourced from literature. Fix well-established values where appropriate to reduce uncertainty.
6. **Rigorous Validation** Conduct thorough validation, including out-of-sample testing, before applying models to decision-making.
7. **Training and Capacity Building** Provide training to ensure managers and researchers can interpret and apply AI-generated models responsibly.

**Figure 9:**
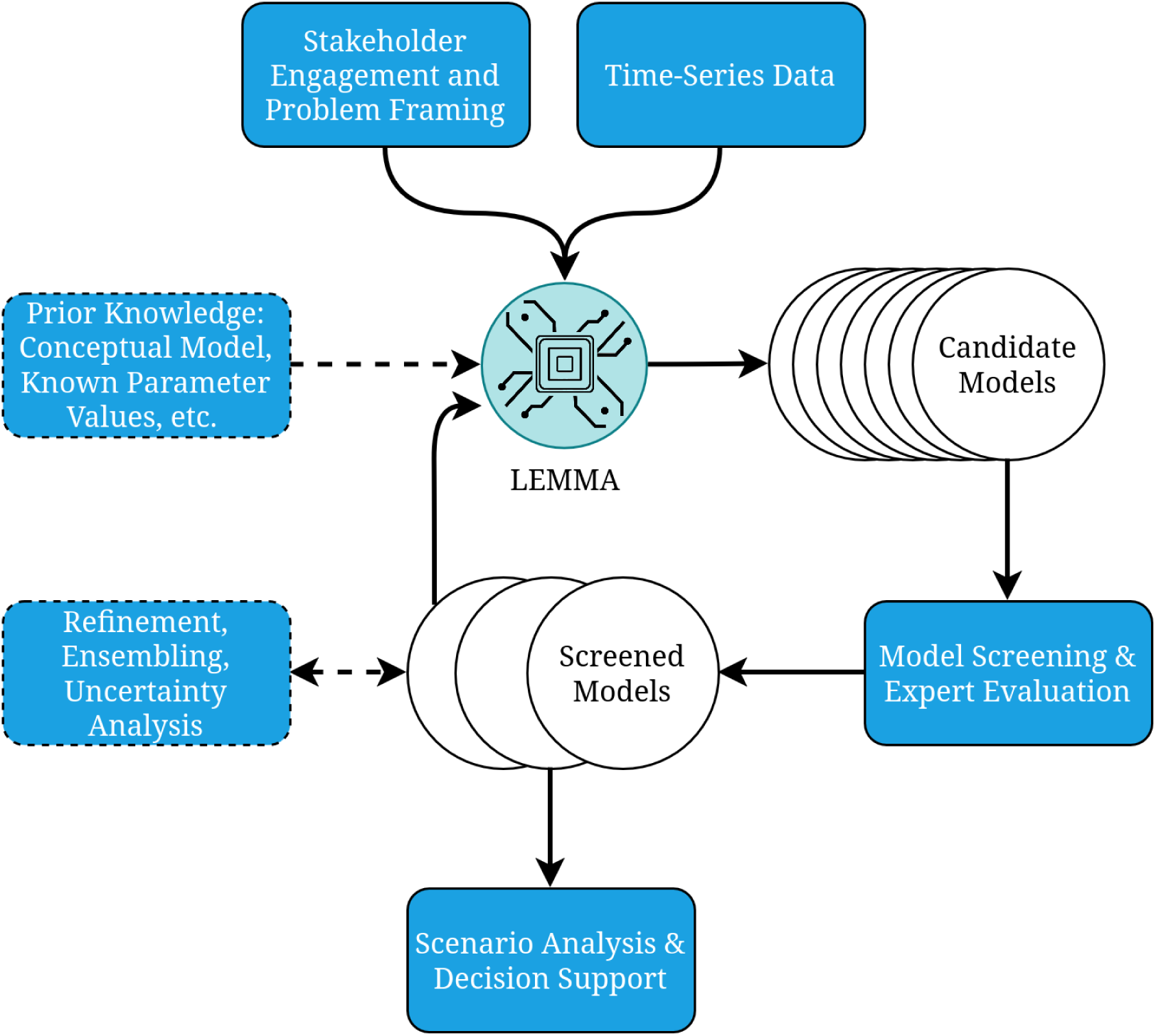
The LEMMA framework workflow integrating human expertise with AI-driven model development. The diagram illustrates how stakeholder engagement, time-series data, and prior ecological knowledge inform the LEMMA process, which generates candidate models that can be evaluated and refined by human experts, ultimately supporting ecosystem management decisions.

In conclusion, LEMMA represents an advancement in ecological modelling that bridges the gap between computational efficiency and ecological insight. By dramatically accelerating model development while maintaining scientific rigour, this framework offers a powerful new tool for researchers and managers facing urgent ecological challenges. As environmental pressures intensify globally, the capacity to rapidly develop, test, and deploy ecologically sound models will become increasingly valuable for effective conservation and management of marine ecosystems.

## Supporting information

Supplemental Results and Methods

## Acknowledgements

SS was supported by an R+ Postdoctoral Fellowship.

## Code and Data Availability

The code for the LEMMA framework is available in a public GitHub repository at manuscriptURL. The repository includes all scripts necessary to reproduce the results presented in this paper, including the genetic algorithm implementation, model evaluation tools, and analysis scripts. The time series data used in the case studies are also available in the repository. The software is released under an MIT license.

## Declaration on Generative AI Usage

During the preparation of this manuscript, we used Claude-3.5-Sonnet and GPT5 to assist with code documentation, manuscript formatting, and language editing. The scientific content, analyses, interpretations, and conclusions presented in this paper were developed and validated by the human authors.

## Author Contributions

SS: Conceptualization, Methodology, Software, Data curation, Formal analysis, Writing - original draft, Writing - review & editing JR: Software, Methodology, Data curation, Writing - review & editing FB: Formal analysis, Supervision, Writing - review & editing BF: Formal analysis, Supervision, Writing - review & editing RT: Supervision, Writing - review & editing MG: Software, Methodology, Writing - review & editing SY: Software, Methodology, Writing - review & editing

## Competing Interests

The authors declare no competing interests.

## Notes

https://github.com/s-spillias/EMs-with-LLMs

### Competing Interest Statement

The authors have declared no competing interest.

### Summary of Updates

This is a revision as a result of the first round of peer-review. Significant changes include: 1. Renaming of the framework to 'LEMMA'. 2. Re-running of full analyses with newer generation of LLMs. 3. Clearer description of methods

