## Supplemental Results and Methods for "Data-Driven Discovery of Mechanistic Ecosystem Models with LLMs"

November 12, 2025

**Keywords:** Artificial Intelligence, Ecological modelling, Evolutionary Algorithms, Large Language Models, Marine Ecosystems, Crown-of-Thorns Starfish, Great Barrier Reef, time-series forecasting

**Abstract**

1. Ecosystem models are essential for ecosystem management, but their development traditionally requires significant time and expertise, creating bottlenecks in addressing urgent environmental challenges.
2. We present ***LEMMA*** (LLM Enabled Mechanistic Modelling for ecosystem Assessment), a framework that programmatically generates and iteratively refines mechanistic ecosystem models by combining large language models (LLMs) for equation synthesis and parameter search, evolutionary algorithms for structural optimization, and Template Model Builder (TMB) for efficient parameter estimation.
3. We critically review *LEMMA*'s ability to recover known ecological relationships through two complementary marine case studies: (1) a nutrient-phytoplankton-zooplankton model, and (2) a Crown-of-Thorns starfish (COTS) model. In the first case, our best models displayed almost perfect recovery of known ecological dynamics while maintaining strong predictive performance across multivariate time-series. In the second case, best *LEMMA* generated models approached human expert models in terms of their ability to successfully capture COTS outbreak dynamics and demonstrated strong out-of-sample predictive power.
4. *LEMMA* produces interpretable models with meaningful parameters that capture real biological processes, facilitating scientific insight and potentially accelerating management applications. By dramatically accelerating model development while offering ecological interpretability, *LEMMA* offers a powerful new tool for addressing urgent ecological challenges in a changing world.

### Data/Code Availability

The data and code associated with this manuscript is available at the anonymised repository: `https://anonymous.4open.science/r/EMs-with-LLMs-10E8`

### 1 Introduction

Ecosystem models provide invaluable information for managing complex interactions between nature and people (McCarthy and Possingham, 2004; Holden and Ellner, 2016), but their development traditionally requires significant time and expertise, creating a bottleneck in addressing urgent environmental challenges (Dichmont et al., 2017; Holden et al., 2024), particularly as climate change demands rapid, adaptable approaches for ecosystem management (Weiskopf et al., 2020; Malhi et al., 2020).

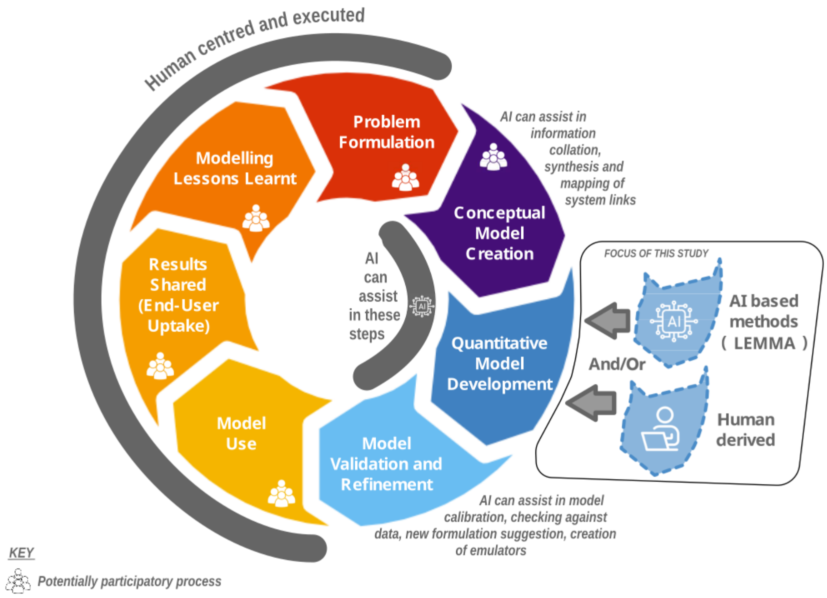


Figure 1: Stylised representation of the iterative modelling process that LEMMA aims to support. Whilst human experts drive the majority of the process, we show that AI-driven processes could play an important role in the Model Development stage.

∞

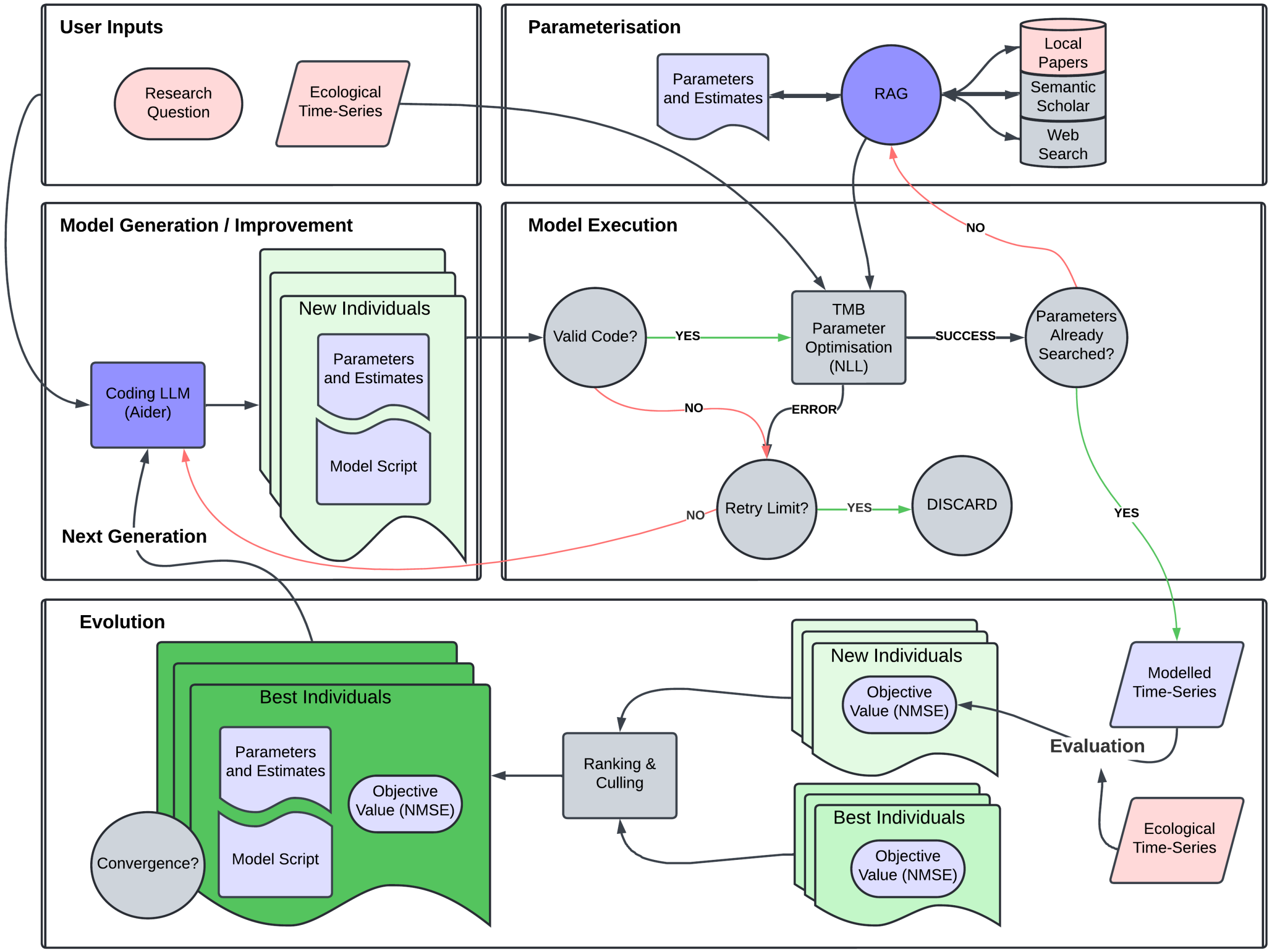


Figure 2: Conceptual diagram of the automated ecological modelling framework, LEMMA. The workflow consists of five main components: (1) User Inputs, where research questions and ecological time-series data are provided; (2) Parameterisation, utilizing RAG-enhanced literature search to estimate parameter values; (3) Model Generation/Improvement, where the Coding LLM creates new individuals with model scripts and parameters; (4) Model Execution, where the LLM's model code is implemented and TMB is used to optimise parameter values; and (5) Evolution, which evaluates model performance through individual assessment, error handling, and ranking-based selection.

##### 2.1.5 Model Evaluation

For each response variable $j$, we calculate a normalized mean squared error:

$$\text{NMSE}_j = \begin{cases} \frac{1}{n}\sum_{i=1}^{n}\left(\frac{y_{ij}-\hat{y}_{ij}}{\sigma_j}\right)^2 & \text{if } \sigma_j \neq 0 \\ \frac{1}{n}\sum_{i=1}^{n}(y_{ij}-\hat{y}_{ij})^2 & \text{if } \sigma_j = 0 \end{cases} \tag{1}$$

where $y_{ij}$ represents observed values for variable $j$ at time $i$, $\hat{y}_{ij}$ represents corresponding model predictions, $\sigma_j$ is the unbiased standard deviation of the observed values for variable $j$ (calculated with $n-1$ denominator), and $n$ is the number of observations. The final objective function value is the mean across all response variables:

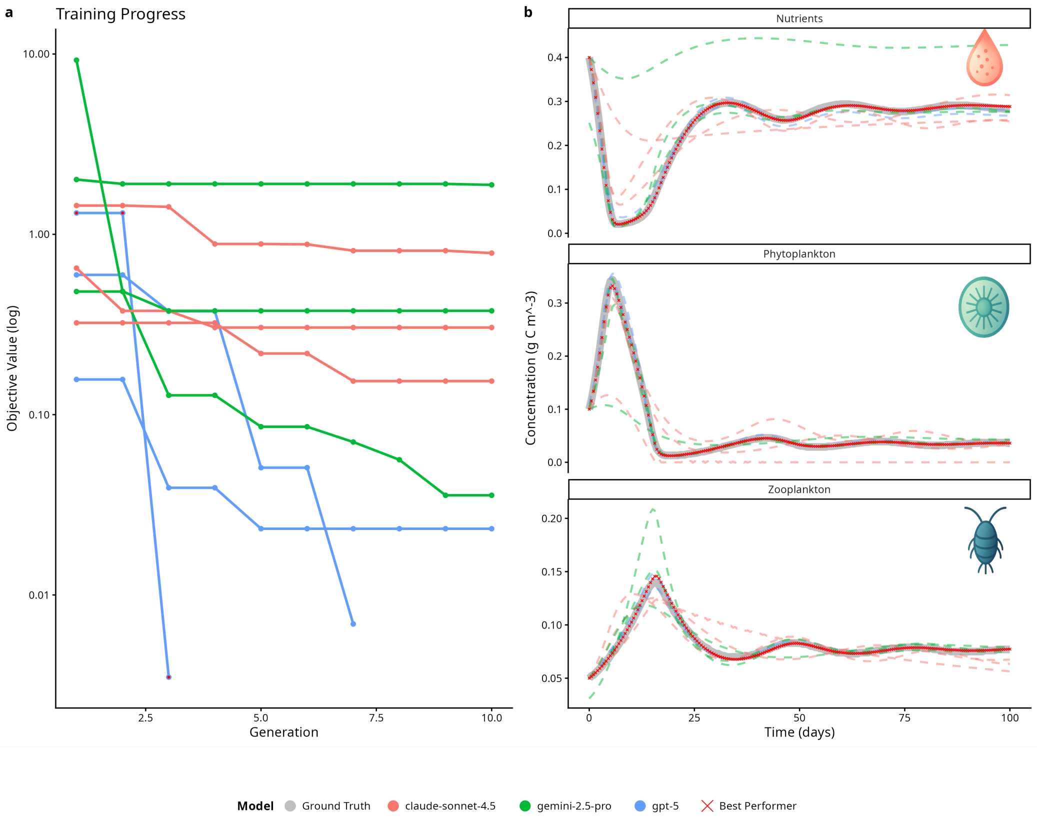


Figure 3: Performance and ecological dynamics of NPZ model retrieval. (a) Training progress showing objective value trajectories across generations (log-scale) for all populations driven by GPT-5, Claude-Sonnet-4.5, and Gemini-2.5-Pro. GPT-5 achieved the lowest objective value (0.004) within three generations. (b) Time-series comparison of nutrient, phytoplankton, and zooplankton concentrations (g C $m^{-3}$) between ground-truth (black lines) and predictions from the best model in each population by objective value (crosses). The model successfully reproduced bloom timing, nutrient draw-down, and trophic phase relationships, indicating strong alignment with ecological mechanisms.

Recovery of NPZ mechanisms varied substantially across LLMs. No model individual LEMMA model perfectly reconstructed all processes, but every mechanism was recovered at least once (Figure 4). GPT-5 achieved the highest overall proportion of well-supported formulations (alternate from Franks (2002) or truth match), with well-supported mechanisms accounting for 80.8% overall versus 19.2% not supported. For the best-performing individual (GPT-5), only phytoplankton mixing was unsupported (not a truth match or alternate). With the exception of both mixing processes, all of the LLMs tended to use supported formulations. Quantitatively, the fluxes of most of the formulations used by the best-performing model were exact or very close matches to those of the truth model. However, the best-perfomer also included a detritus pathway that was not present in the ground-truth (Figure 5). These patterns indicate that while overall ecological fidelity was high, certain processes, like phytoplankton and nutrient mixing were often difficult (but not impossible) for LLMs to reconstruct accurately and that quantitative matches were substantiated by ecological intention.

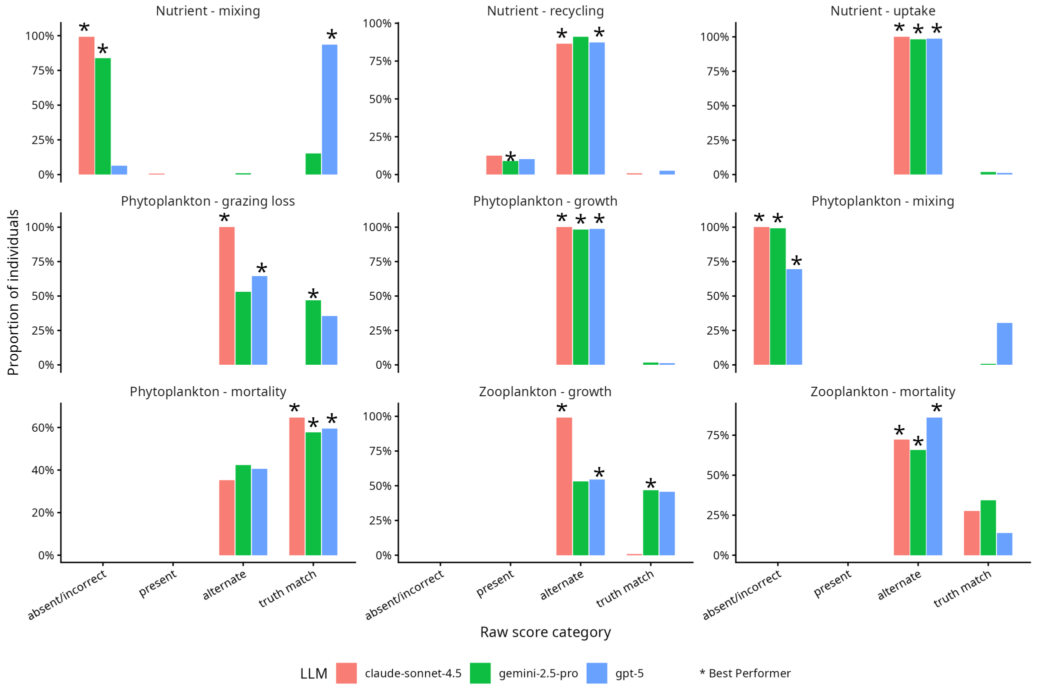


Figure 4: Mechanism-level recovery of NPZ equations across LLM families. Each panel shows the distribution of raw scores for a specific ecological mechanism as described in Section 2.2.2

In addition to the presence of the ground-truth processes, we also saw that the best-performing LEMMA model successfully reproduced most ecological dynamics of the NPZ system according to the specific fluxes, although it also included additional processes that were not present in the ground truth model (Figure 5). Uptake, phytoplankton growth, and zooplankton mortality exhibited near-perfect alignment with the ground-truth model, indicating that core trophic interactions were accurately reconstructed. Grazing loss and recycling processes were also well captured, though minor deviations were observed in recycling to nitrogen at higher flux values. As reported above, phytoplankton mixing was absent in the LEMMA-derived model. The best-performing model also included a detritus pathway which was not present in the original ground-truth model.

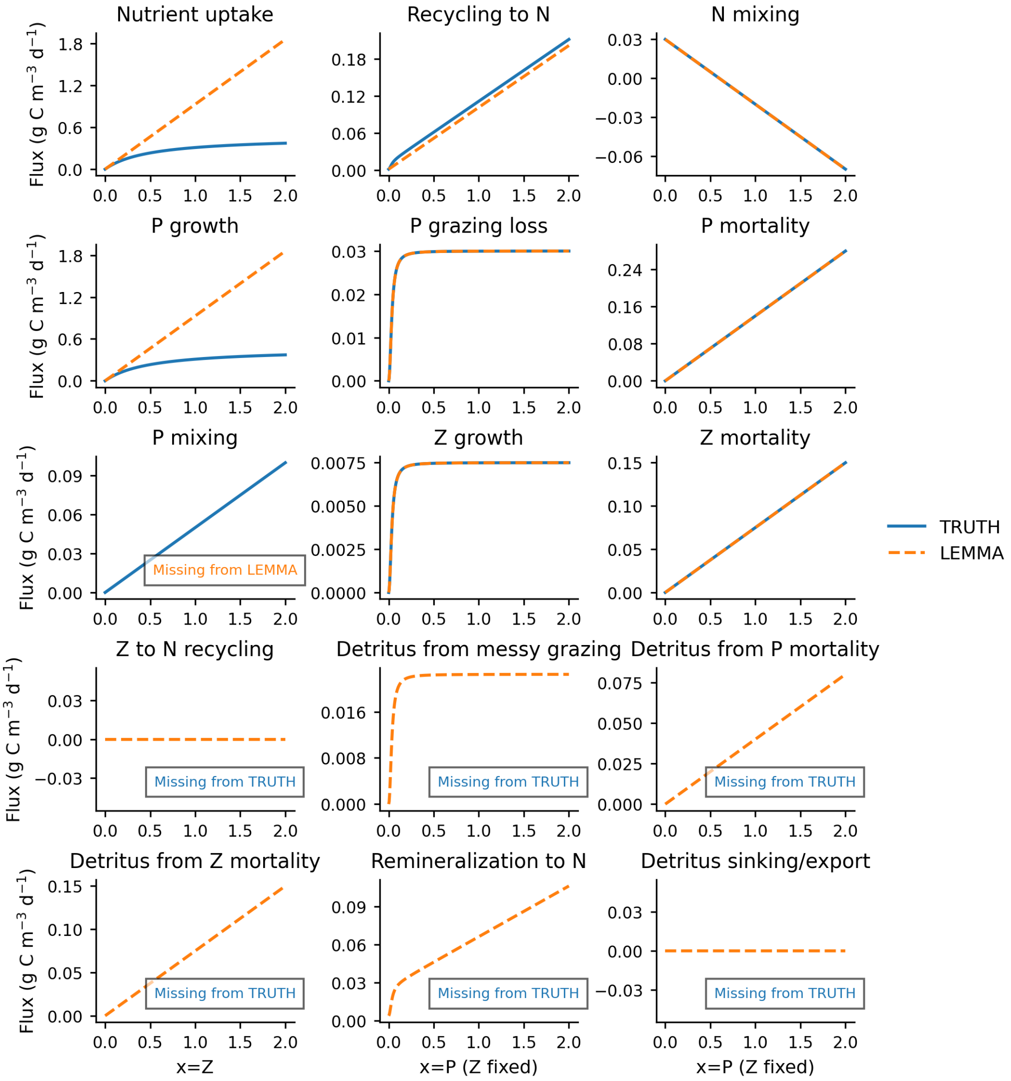


Figure 5: Comparison of process-level fluxes between the ground-truth NPZ model and best-performing LEMMA-derived model. Each panel shows the functional form of a single flux term plotted against its primary driver, with other state variables held constant at representative initial values. Optional modifiers in the compiled model (temperature, light, quadratic zooplankton mortality, detrital recycling) are disabled to isolate core equation differences. Parameter values are aligned so that curve differences reflect structural form rather than parameterization. Processes absent in one model are indicated as missing.

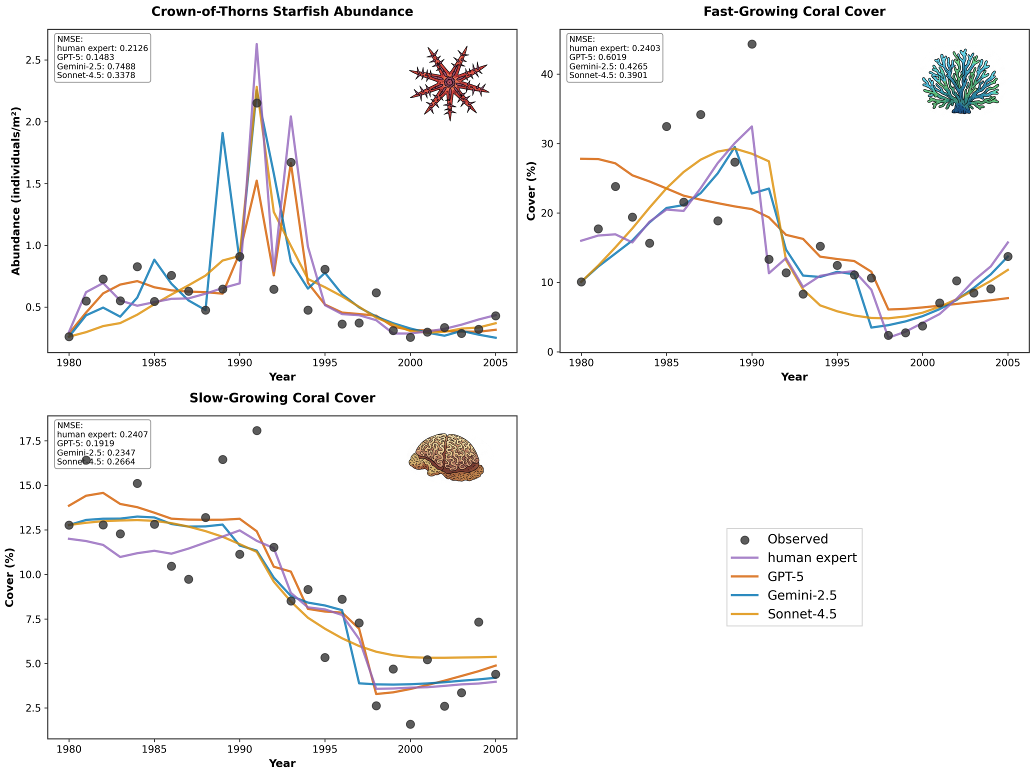


Figure 6: Comparison of model predictions across ecosystem components. The plots show observed versus predicted values for COTS abundance, fast-growing coral cover, and slow-growing coral cover, demonstrating the models' ability to capture key ecological patterns and relationships. Objective values (obj) shown in the legend represent the normalised mean squared error for all three variables, where lower values indicate better model performance.

##### 3.3.1 Temporal Hold-Out

Because GPT-5 demonstrated the strongest overall performance, we used it as the base LLM to test whether the LEMMA framework could create models capable of predicting out-of-sample datapoints. After ten generations, we found that the best-performing model (objective value: 0.29) had a reasonable degree of out-of-sample prediction performance on the withheld test dataset, with particularly strong predictive power for fast-growing coral cover ($R^2$ = 0.81, RMSE = 2.22, MAE = 1.92). Interestingly, this was a better objective value than the previous experiments where LEMMA had access to the entire time-series.

For slow-growing coral cover, the model achieved moderate predictive accuracy ($R^2 = 0.17$, RMSE = 1.6, MAE = 1.3), effectively capturing the general declining trend while showing some deviation in precise values. COTS population predictions demonstrated strong accuracy ($R^2 = 0.75$, RMSE = 0.07, MAE = 0.05), successfully capturing the timing and magnitude of major outbreaks.

Figure 7 illustrates these prediction capabilities, showing both training period performance (pre-1997) and out-of-sample predictions (1997-2005). The model's ability to maintain consistent error metrics (RMSE and MAE) while capturing both rapid population dynamics and slower coral cover changes suggests it successfully identified fundamental ecological relationships governing this system.

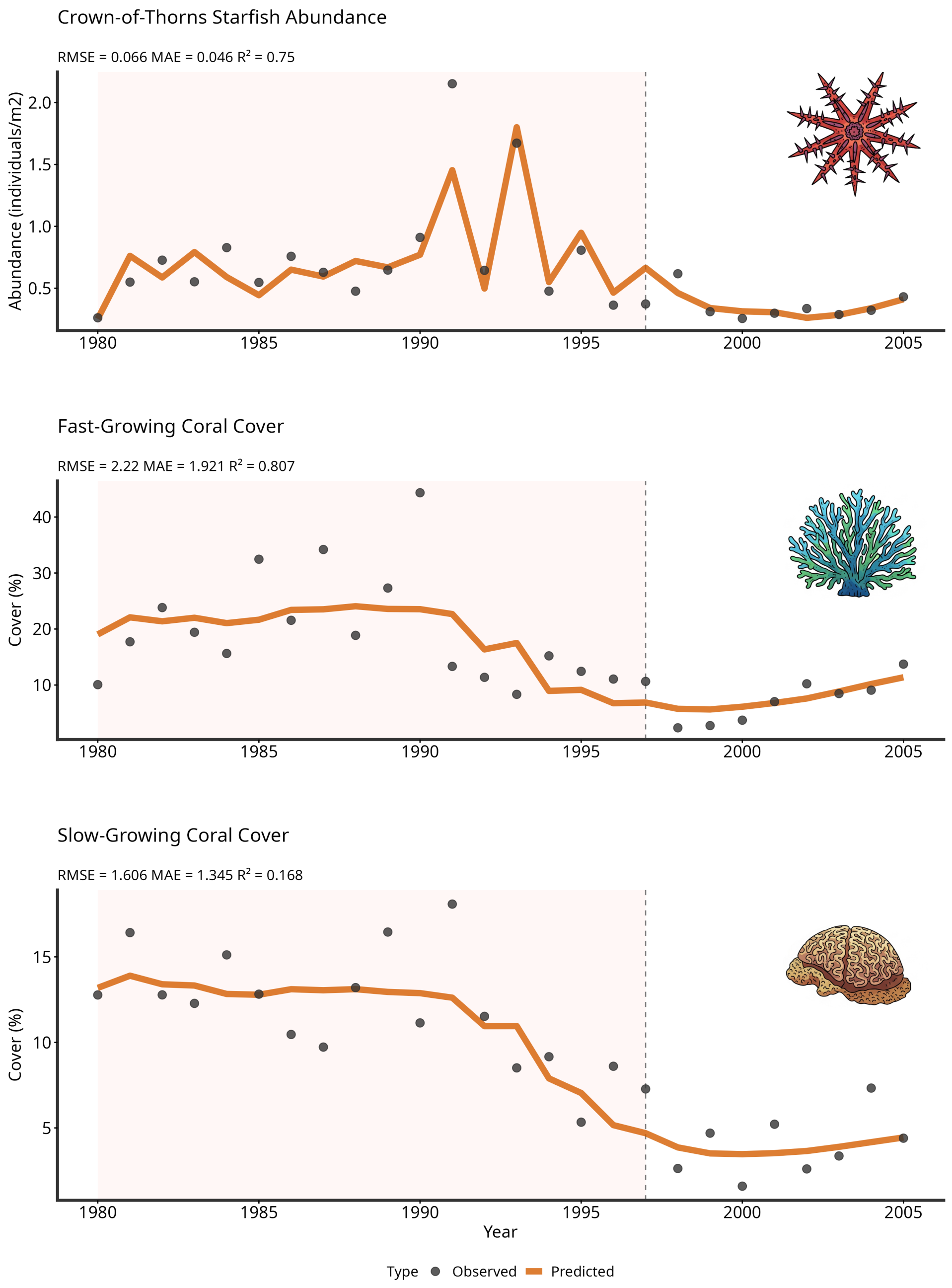


Figure 7: Temporal hold-out evaluation of LEMMA driven by the best-performing LLM (GPT-5) showing predictions against observed data. LEMMA was provided with 70% of the time-series data (pink shaded region) and developed a model that was then evaluated on the full time-series, including the remaining unseen 30% of the time-series data (white region). Lines represent model predictions, dots show observed data.

#### 3.4 Parameter Retrieval

We assessed how frequently LEMMA linked model parameters to external evidence sources, including Semantic Scholar searches and a locally curated document store. Across all generated models (21 populations, 2,619 parameters), citation integration was modest: only 3.1% of parameters were linked to explicit citations, 1.2% to Semantic Scholar matches, and 2.0% to document store references. The majority of parameters lacked any external linkage. Differences emerged between case studies. NPZ-based populations showed slightly higher citation rates (3.7%) and semantic matches (2.7%) than COTS-focused populations (2.8% and 0.59%, respectively). Document store usage was notably higher for COTS models (2.4%) compared to NPZ (1.1%). This disparity likely reflects the fact that the document store was curated specifically for the COTS case study, whereas no curation was performed for NPZ, limiting retrieval opportunities. At the individual model level, citation coverage rarely exceeded 10% of parameters, and many individuals contained no cited parameters. Where citations were present, they typically supported temperature optima, growth coefficients, and bleaching thresholds, parameters with well-documented ecological relevance. These results indicate that while LEMMA can incorporate literature-based evidence, this capability was underutilized in practice. Enhancing citation integration may require prioritizing literature-grounded parameter selection and expanding curated resources for all ecosystem contexts.

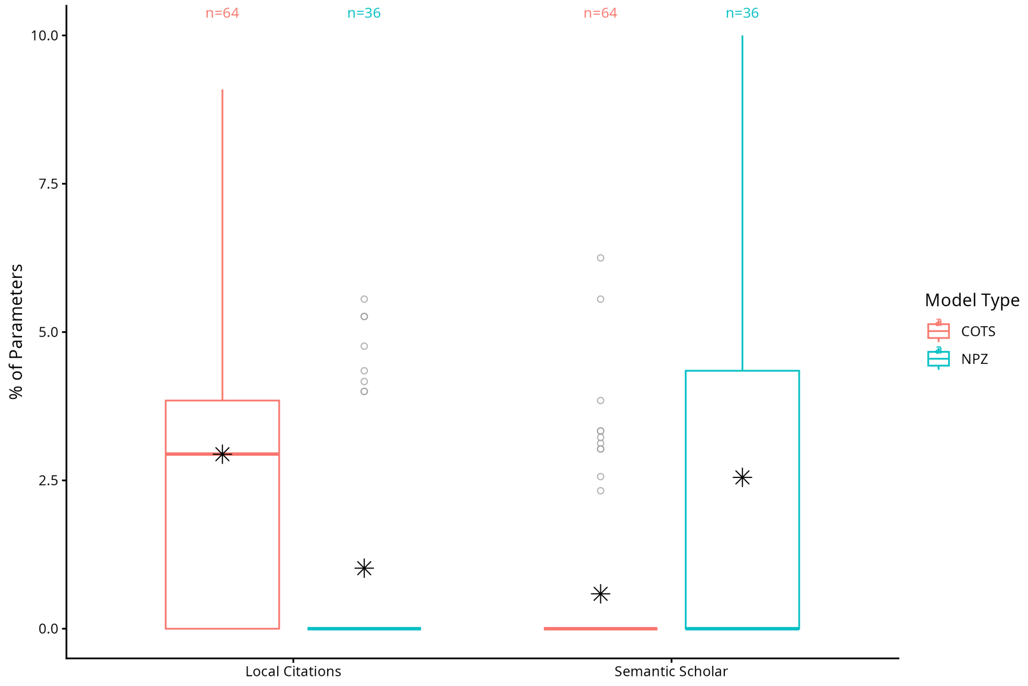


Figure 8: Comparison of citation integration between NPZ and COTS case studies. Boxplots illustrate the proportion of model parameters with citations (Semantic Scholar or local document store). NPZ models were more reliant on Semantic Scholar because the local document store was curated specifically for the COTS case study. Asterisks represent the best performer (lowest nRMSE objective value) for each model type.

2. **Expert Review** All AI-generated models should be reviewed by do-

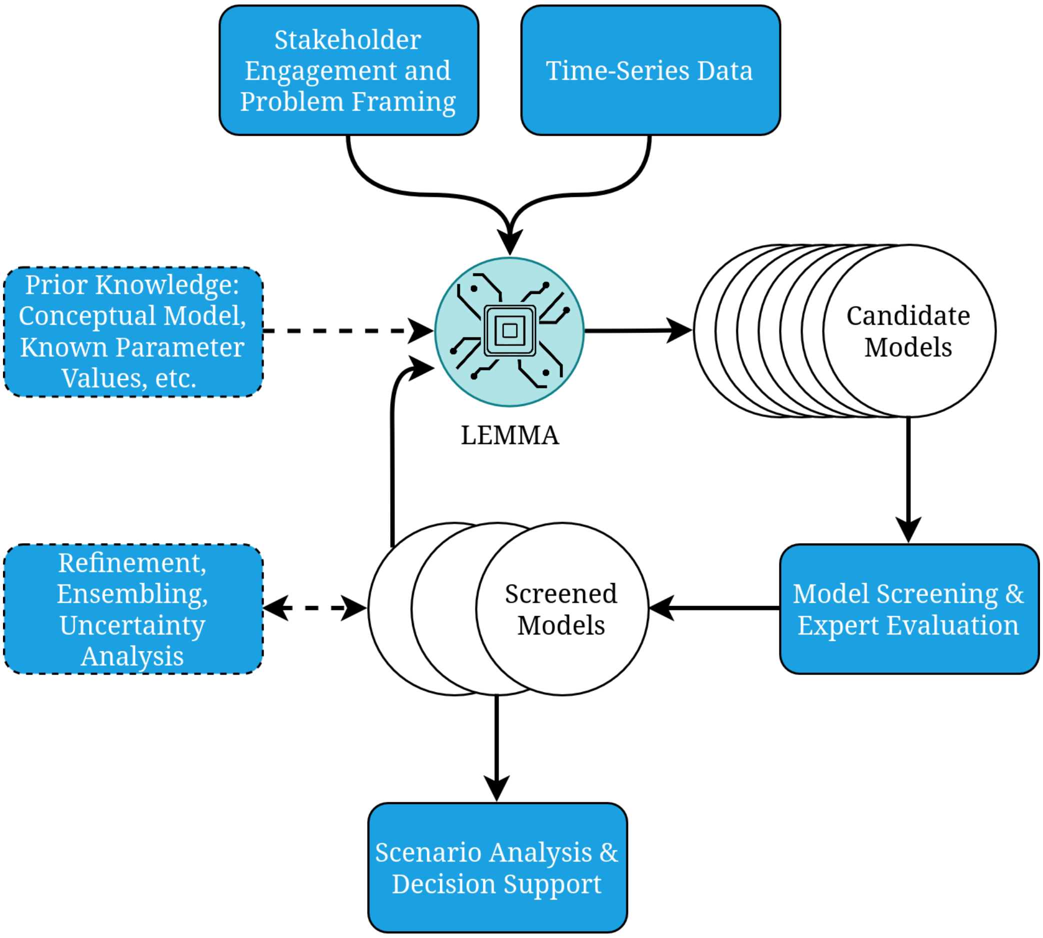


Figure 9: The LEMMA framework workflow integrating human expertise with AI-driven model development. The diagram illustrates how stakeholder engagement, time-series data, and prior ecological knowledge inform the LEMMA process, which generates candidate models that can be evaluated and refined by human experts, ultimately supporting ecosystem management decisions.

main experts to validate ecological plausibility and ensure alignment with management objectives.

3. **Complementary Use** Use LEMMA as a complementary tool that supports - rather than replaces - traditional modelling workflows, especially for rapid prototyping or exploring alternative hypotheses.

4. **Model Transparency** Maintain transparency through clear documentation of equations, parameters, and assumptions to ensure traceability and reproducibility.

5. **Parameter Assessment** Critically assess parameter values sourced from literature. Fix well-established values where appropriate to reduce uncertainty.

6. **Rigorous Validation** Conduct thorough validation, including out-of-sample testing, before applying models to decision-making.

7. **Training and Capacity Building** Provide training to ensure managers and researchers can interpret and apply AI-generated models responsibly.

In conclusion, LEMMA represents an advancement in ecological modelling that bridges the gap between computational efficiency and ecological insight. By dramatically accelerating model development while maintaining scientific rigour, this framework offers a powerful new tool for researchers and managers facing urgent ecological challenges. As environmental pressures intensify globally, the capacity to rapidly develop, test, and deploy ecologically sound models will become increasingly valuable for effective conservation and management of marine ecosystems.
